# Neuronal Activity-Induced BRG1 Phosphorylation Regulates Enhancer Activation

**DOI:** 10.1101/2020.09.01.278101

**Authors:** Bongwoo Kim, Yi Luo, Xiaoming Zhan, Zilai Zhang, Xuanming Shi, Jiaqing Yi, Zhenyu Xuan, Jiang Wu

**Author notes:** These authors contributed equally and are listed alphabetically (Bongwoo Kim, Yi Luo). **CORRESPONDENCE:** Jiang Wu, PhD, Department of Physiology, UT Southwestern Medical Center, 5323 Harry Hines Blvd. Dallas TX 75390-9040.

## Abstract

Neuronal activity-induced enhancers drive the gene induction in response to stimulation. Here, we demonstrate that BRG1, the core subunit of SWI/SNF-like BAF ATP-dependent chromatin remodeling complexes, regulates neuronal activity-induced enhancers. Upon stimulation, BRG1 is recruited to enhancers in an H3K27Ac-dependent manner. BRG1 regulates enhancer basal activities and inducibility by affecting cohesin binding, enhancer-promoter looping, RNA polymerase II recruitment, and enhancer RNA expression. Furthermore, we identified a serine phosphorylation site in BRG1 that is induced by neuronal activities and is sensitive to CaMKII inhibition. BRG1 phosphorylation affects its interaction with several transcription co-factors, possibly modulating BRG1 mediated transcription outcomes. Using mice with knock-in mutations, we showed that non-phosphorylatable BRG1 fails to efficiently induce activity-dependent genes, whereas phosphomimic BRG1 increases the enhancer activities and inducibility. These mutant mice displayed anxiety-like phenotypes and altered responses to stress. Therefore, our data reveal a mechanism connecting neuronal signaling to enhancer activities through BRG1 phosphorylation.

## INTRODUCTION

The abilities of neurons to respond to various stimulations and to convert transient stimuli into long-term changes in brain function underlies neuron reactivity and plasticity. Activity-regulated gene (ARG) expression plays a central role in short-term neural responses as well as in long-term memory formation, homeostasis, and adaptation (Ebert and Greenberg, 2013; Ganguly and Poo, 2013; West and Greenberg, 2011; Yap and Greenberg, 2018). ARGs include immediate early genes (IEGs) that are induced within minutes and late response genes (LRGs) that are induced over hours (Tyssowski et al., 2018; Yap and Greenberg, 2018). Some IEGs encoding transcription factors such as c-FOS and NPAS4 regulate the immediate stress response and may mediate the expression of LRGs (Brigidi et al., 2019; Malik et al., 2014). The activity-induced expression of ARGs such as *Arc, Bdnf*, and *Igf1* are important for synaptic maturation and homeostasis in various neuron subtypes (Hong et al., 2008; Jakkamsetti et al., 2013; Mardinly et al., 2016). Altered expression of ARGs cause changes in synaptic activity, neuron morphology, and circuit formation that may lead to behavior defects in stress responses, learning and memory, addiction, and psychiatric disorders (Clayton et al., 2019; Gallo et al., 2018; Manning et al., 2017; Yap and Greenberg, 2018).

ARG expression is directed through activity-induced enhancers by specific neuronal stimuli (Joo et al., 2016; Kim et al., 2010; Malik et al., 2014; Nord and West, 2020; Tyssowski et al., 2018). Activated distal enhancers that are marked by H3K27Ac interact with promoters to form enhancer-promoter looping (Kim and Shiekhattar, 2015). Cohesin-mediated enhancer-promoter looping facilitates RNA polymerase II (RNA Pol II) recruitment and enhancer RNA (eRNA) expression, which then drives mRNA transcription initiation and elongation (Kagey et al., 2010; Kim and Shiekhattar, 2015; Ong and Corces, 2011). Thousands of neuronal activity-induced enhancers have been identified in cultured primary cortical neurons in response to KCl-mediated depolarization (Kim et al., 2010; Malik et al., 2014). These enhancers are marked by increased H3K27Ac and overlap with enhancers identified in developing mouse brains (Shen et al., 2012). Inducible enhancers connect various signals to gene expression through the cooperation between transcription factors and chromatin regulators. Chromatin states could be instructive or permissive for gene expression through regulation of DNA and histone accessibility to transcription factors and the general transcription machinery (Allis and Jenuwein, 2016; Talbert et al., 2019). Many chromatin regulators, such as histone acetyl transferase CBP and histone deacetylases (HDACs), methyl DNA recognizing protein MECP2, and chromatin remodeling regulators BAF and NuRD complexes, have been shown to play important roles in ARG regulation (Chen et al., 2019; Chen et al., 2003; Qiu and Ghosh, 2008b; Yang et al., 2016; Zhang et al., 2016). How these factors respond to neural signaling and act on enhancers are under intense study. Importantly, it remains largely unclear how they functionally interact with each other to modulate transcription outcomes.

Mammalian SWI/SNF-like ATP-dependent chromatin remodeling BAF complexes, which contain core ATPase subunits BRG1 (also known as SMARCA4) or BRM (also known as SMARCA2) and 10-12 tightly associated subunits, use energy derived from ATP hydrolysis to modulate chromatin structures and regulate transcription (Kasten et al., 2011; Son and Crabtree, 2014; Wu, 2012). Mutations in BAF subunits are the genetic causes of Coffin-Siris syndrome (Santen et al., 2012; Tsurusaki et al., 2012; Van Houdt et al., 2012), which results in severe neural developmental defects. In addition, *de novo* functional mutations in genes encoding BAF subunits are observed in patients with autism spectrum disorders, amyotrophic lateral sclerosis, and schizophrenia (De Rubeis et al., 2014; Halgren et al., 2012; Helsmoortel et al., 2014; Neale et al., 2012). Moreover, *BRG1* is a key node in the autism spectrum disorder gene network (De Rubeis et al., 2014). Animal studies demonstrated important functions of BAF subunits in neural development and plasticity (Sokpor et al., 2017; Son and Crabtree, 2014; Wu, 2012).

BRG1 and BAF complexes preferentially bind to enhancers during development and during cancer development and progression (Alexander et al., 2015; Alver et al., 2017; Yu et al., 2013). The biochemical and molecular mechanisms that regulate BAF function at enhancers remain largely uncharacterized, however. Mechanistic analyses are complicated because BAF complexes can function as activators or as repressors depending on the subunit composition, chromatin environment, and interacting epigenetic regulators. Previously, we and others have shown that neuronal BAF complexes are important for ARG expression and synaptic development and plasticity (Qiu and Ghosh, 2008b; Wu et al., 2007; Zhang et al., 2016). The rapid and relatively synchronized neuronal ARG gene induction in response to neuronal stimulation provides an ideal platform to directly dissect BAF function in enhancer activation.

Neuronal activities, which trigger Ca^2+^ influx through voltage-sensitive calcium channels and N-methyl-D-aspartate receptors, initiate multiple signaling pathways mediated by intermediates, often kinases or phosphatases, that transduce signals into the nucleus to regulate transcription (Deisseroth et al., 2003; Ebert and Greenberg, 2013; Greer and Greenberg, 2008; Qiu and Ghosh, 2008a). These intermediates, such as cAMP/PKA, Ras/MAPK, CaMK, and calcineurin, can phosphorylate or dephosphorylate a number of transcription factors and co-factors that serve as activity switches to regulate ARG expression (Ebert and Greenberg, 2013; Wong and Ghosh, 2002; Yap and Greenberg, 2018). Phosphorylation of transcription regulators influences a dynamic protein interaction network that allows neurons to produce rapid and diverse responses to enable adaption to the changing environment. Despite the identification of phosphorylation sites in BAF subunits in various conditions (Kimura et al., 2014; Kwon et al., 2015; Padilla-Benavides et al., 2020; Wang et al., 2013), it is not clear whether BAF subunits undergo protein modifications in response to neuron activities or how post-translational modifications regulate BAF activities.

To characterize the function of BRG1 in ARG regulation in response to neuronal activities, we used a combination of genetic, genomic, and proteomic approaches. We demonstrated that BRG1 is recruited to enhancers upon neuronal stimulation in an H3K27Ac-dependent manner. We showed that BRG1 regulates enhancer activities by affecting cohesin binding, enhancer-promoter looping, RNA pol II recruitment, and eRNA expression. We further identified a dynamic serine phosphorylation site in BRG1 that is induced by neuronal activities. BRG1 phosphorylation affects its interaction with several key transcription co-factors, which may underlie the BRG1-regulated ARG transcription outcomes. Using knock-in mice generated using the CRISPR-Cas9 approach, we showed that phosphorylation modulates BRG1 activities in ARG enhancer activation. Mice that express a non-phosphorylatable BRG1 or a phosphomimic BRG1 displayed anxiety-like phenotypes and altered responses to stress. Therefore, we uncovered a mechanism underlying BRG1 function in regulating enhancer activities and identified a novel phosphorylation event that fine-tunes BRG1 function in response to neuronal signaling. Our study provides significant insights into the function of chromatin remodeling complexes in neural development and plasticity.

## RESULTS

### BRG1 regulates neuronal ARG activation

Using a mouse pan neuron Cre line (*BAF53b-Cre*), we specifically deleted *Brg1* in all developing neurons resulting in the conditional *Brg1* knock-out line (*Brg1^cko^* mice) (Zhan et al., 2015; Zhang et al., 2016). Previously, we showed that in cultured *Brg1^cko^* cortical neurons, *Brg1* deletion, which happened gradually over several days in culture, specifically affected the expression of a number of ARGs such as *Bdnf* and *Nr4a1* at 6 hours after KCl-mediated depolarization (Zhang et al., 2016). The analyses of the previous RNA-seq results (Zhang et al., 2016) showed that 76 ARGs displayed increased gene expression in wild-type neurons after KCl treatment, but had reduced induction levels in *Brg1^cko^* neurons (Figure 1A). The analyses at 6-hour time point may miss some BRG1-regulated IEGs. A more detailed time course study of neuronal gene expression after KCl-induced depolarization showed that *Brg1* deletion led to impaired expression of not only LRGs such as *Bdnf* but also of IEGs such as *c-Fos* and *Arc* (Figure 1B, S1). Although many ARGs were affected by *Brg1* deletion, several genes, such as *Gadd45b* (Figure S1) and *Junb*, were not (Zhang et al., 2016), indicating functional specificities of BRG1 and the overall normal Ca^2+^ signaling in response to depolarization. These results demonstrated that BRG1 is critical for ARG expression.

**Figure 1.**
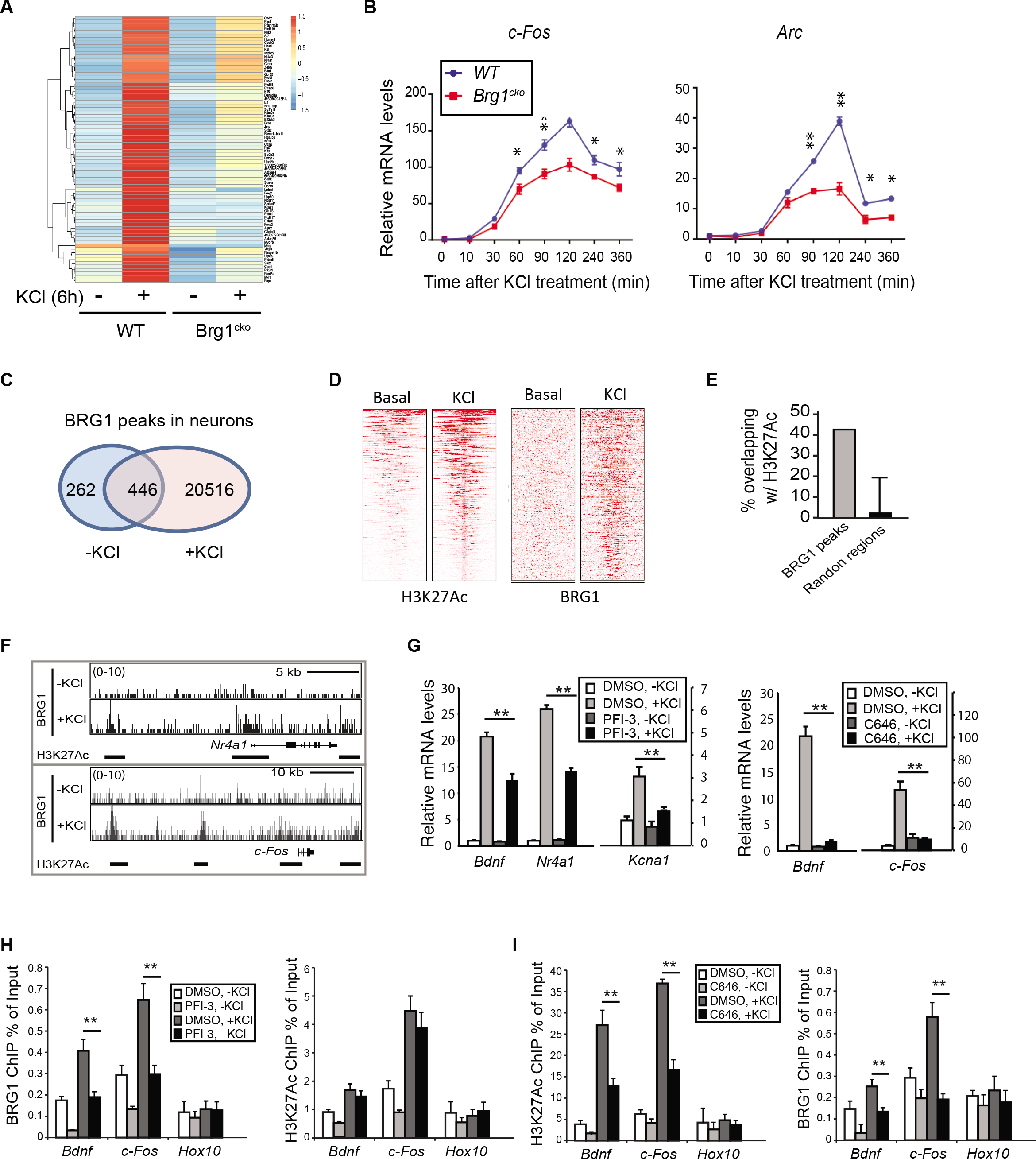
BRG1 regulates ARG induction and binds to H3K27Ac marked enhancers in response to neuronal activities. A. Heatmap showing RNA-seq signals of 76 ARGs that were differentially expressed in *Brg1^cko^* neurons compared to wild-type neurons at 6 h after KCl treatment. There were no significant differences of their expression in resting wild-type and *Brg1^cko^* neurons. B. Relative mRNA levels of *c-Fos* and *Arc* in cultured *Brg1^cko^* (red square) and control (blue dot) neurons after KCl treatment as shown by RT-qPCR. C. Overlaps between BRG1 ChIP-seq peaks under basal and I h KCl-stimulated conditions in cultured cortical neurons. D. Heatmap of H3K27Ac and BRG1 ChIP-seq signals in 10-kb regions surrounding all H3K27Ac peaks that showed increased signals upon depolarization in cultured cortical neurons. E. Percent overlap between BRG1 peaks and randomly selected regions with H3K27Ac peaks in depolarized neurons. F. BRG1 ChIP-seq signals in resting and depolarized neurons around activity-induced genes *Nr4a1* and *c-Fos*. H3K27Ac peaks in depolarized neurons are shown as bars below the BRG1 ChIP-seq plot. G. Relative mRNA levels determined by RT-qPCR in cultured cortical neurons treated with BAF bromodomain inhibitor PFI-3 or CBP/p300 inhibitor C646 with or without a 6-h KCl treatment. H. ChIP-qPCR assays of BRG1 and H3K27Ac levels in cultured cortical neurons treated with or without PFI-3 and with or without 0.5-h KCl treatment. I. ChIP-qPCR assays of BRG1 and H3K27Ac levels in cultured cortical neurons treated with or without C646 and with or without 0.5-h KCl treatment. Significance was determined by Student’s t-test (n=3). *: p<0.05. **: p<0.01.

### Activity-induced BRG1 binding to enhancers requires H3K27Ac

BAF complexes may regulate ARG expression through direct chromatin binding. To determine where BRG1 binds on chromatin and how binding changes in response to neuronal activities, we performed BRG1 ChIP-seq in resting and KCl-depolarized cortical neuron cultures (50 mM KCl for 1 h). We found that induction of neuronal activity dramatically increased BRG1 binding: peak numbers increased from 708 to 20,962 (Figure 1C). A previous genome-wide study revealed that dynamic H3K27Ac modification marks were also enhanced by neuron depolarization in a similar experimental setting (Kim et al., 2010; Malik et al., 2014). An intersection between BRG1 and H3K27Ac peaks showed that most activity-induced H3K27Ac sites also had increased BRG1 binding upon depolarization (Figure 1D). A distribution analysis showed that over 40% of the BRG1 peaks overlapped with H3K27Ac peaks in depolarized neurons (Figure 1E). Examination of the regulatory regions of ARGs such as *c-Fos* and *Nr4a1* showed extensive co-occupancies by BRG1 and H3K27Ac in enhancer peaks in depolarized neurons (Figure 1F). This observation is consistent with previous observations of BAF enrichment at enhancers during development and in cancer models (Alexander et al., 2015; Alver et al., 2017; Hodges et al., 2018).

The BAF chromatin remodeling complex could be recruited to specific chromatin sites through interactions with transcription factors that bind sequence specifically or through interactions with histones (Wu et al., 2009). Transcription factors such as MEF2, SP1, and AP-1 have been shown to be important for BRG1 recruitment to ARG regulatory regions (Qiu and Ghosh, 2008b; Vierbuchen et al., 2017; Zhang et al., 2016). In addition, the bromodomain of BRG1 binds to H3K27Ac with low affinity (Filippakopoulos et al., 2012). Therefore, BAF complexes could also be recruited to enhancers through interactions with H3K27Ac, which is rapidly increased upon depolarization (Malik et al., 2014). To determine whether the BRG1 bromodomain is important for BAF recruitment to H3K27Ac-marked active enhancers, we treated cultured neurons with a BRG1 bromodomain inhibitor PFI-3 (Gerstenberger et al., 2016) or with C646, which is an inhibitor of histone acetyltransferase (HAT) p300/CBP (Bowers et al., 2010) before KCl treatment. Both PFI-3 and C646 significantly impaired activity-induced BRG1 target gene expression (Figure 1G). Importantly, ChIP-qPCR showed that PFI-3 treatment impaired activity-induced BRG1 binding to enhancers but did not significantly affect activity-induced local H3K27Ac levels shortly after depolarization (Figure 1H). In contrast, C646 treatment not only reduced local H3K27Ac levels but also impaired activity-induced BRG1 binding to the enhancers (Figure 1I). These data suggest that at early stage of ARG induction, H3K27Ac helps recruit BRG1 to activity-induced enhancers through an interaction with the BRG1 bromodomain.

### BRG1 regulates activity-induced enhancer-promoter looping and enhancer activities

We showed that *Brg1* deletion led to impaired induction of ARGs in response to depolarization. Consistent with this finding, we observed a reduction in RNA polymerase II (RNA pol II) recruitment to the promoters of several ARGs (Figure 2A). Since BRG1 binds to enhancers, we examined the levels of the active enhancer marker H3K27Ac in wild-type and *Brg1^cko^* neurons. *Brg1* deletion reduced H3K27Ac levels at several well characterized activity-induced enhancers (Figure 2B) including the *c-Fos* enhancer 2 and the *Arc* enhancer (Schaukowitch et al., 2014). Therefore, although the initial rapid increase of H3K27Ac is required for BRG1 recruitment, BRG1 is also required for later maximum induction of H3K27Ac. Thus, epigenetic regulators, including both the BAF chromatin remodeling complex and p300/CBP H3K27 HATs, work cooperatively to activate ARGs. Since eRNA expression levels often correlate with enhancer activities and gene expression, we examined the levels of eRNAs previously reported to influence *c-Fos* and *Arc* expression (Schaukowitch et al., 2014). We observed defects in activity-induced expression of these eRNAs upon *Brg1* deletion (Figure 2C).

**Figure 2.**
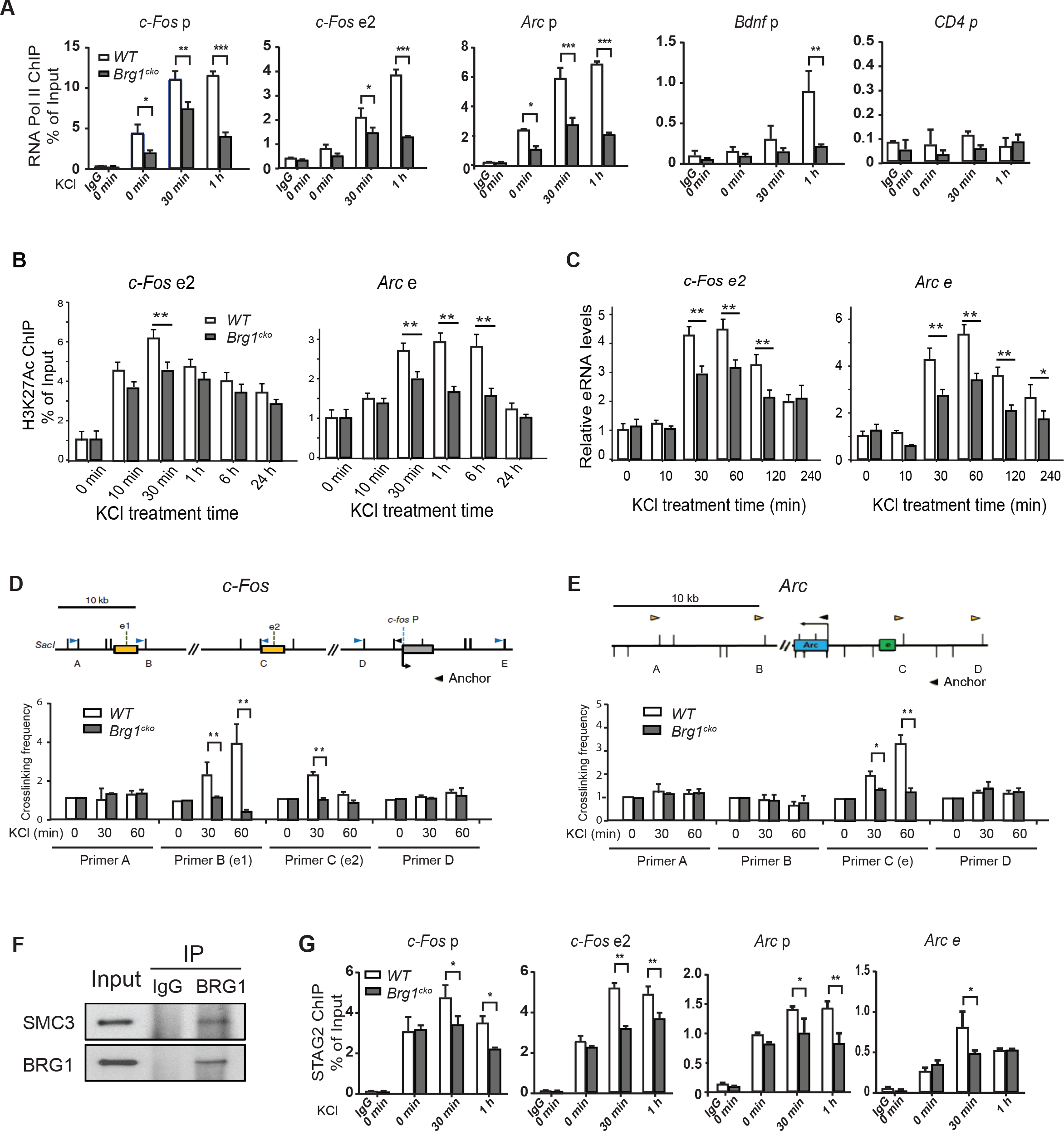
BRG1 regulates activity-induced enhancer-promoter looping and enhancer activities. A. ChIP-qPCR assay of RNA Pol II binding to the promoters and enhancers of ARGs in wild-type and *Brg1^cko^* neurons at indicated times after KCl stimulation. The *CD4* gene was shown as a negative control. B. H3K27Ac levels at *c-Fos* enhancer 2 (e2) and *Arc* enhancer (e) in wild-type and *Brg1^cko^* neurons at indicated times after KCl stimulation measured by ChIP-qPCR. C. Relative eRNA levels of *c-Fos* enhancer 2 and *Arc* enhancer in *Brg1^cko^* and control neurons after KCl treatment as shown by RT-qPCR. D. Crosslinking frequency in 3C experiments between enhancers and promoter of the *c-Fos* gene in basal or depolarized wild-type and *Brg1^cko^* neurons. The genomic structures and the PCR primer locations are shown above the results. The anchor primer was paired with primer B or C for measuring the interactions between *c-Fos* promoter and enhancer 1 or enhancer 2, respectively. Primer A and D were used for negative control regions. E. Crosslinking frequency in 3C experiments between enhancers and promoter of the *Arc* gene in basal or depolarized wild-type and *Brg1^cko^* neurons. The genomic structures and the PCR primer locations are shown above the results. The anchor primer was paired with primer C for measuring the interactions between *Arc* promoter and enhancer. Primer A, B, and D were used for negative control regions. F. Western blot analysis of endogenous SMC3 co-immunoprecipitated with BRG1 from post-natal day 5 (P5) cortex tissues. G. ChIP-qPCR analysis of STAG2 levels at enhancers and promoters of *c-Fos* and *Arc* genes in wild-type and *Brg1^cko^* neurons at indicated times after KCl stimulation. Significance was determined by Student’s t-test (n=3). *: p<0.05. **: p<0.01.

To determine the direct function of BRG1 at enhancers, we examined an early critical step of enhancer activation, enhancer-promoter looping. Using the chromatin conformation capture (3C) assay, we examined the function of BRG1 in enhancer-promoter looping in *c-Fos* and *Arc* gene regulatory regions. It was previously shown that both enhancer 1 and enhancer 2 of the *c-Fos* gene have increased levels of contact with the *c-Fos* promoter in response to neuronal depolarization (Joo et al., 2016). We confirmed the activity-induced looping of these enhancer-promoter pairs in wild-type neurons; whereas interactions between the promoter and other regions were not increased by KCl treatment (Figure 2D). *Brg1* deletion significantly impaired the *c-Fos* enhancer-promoter looping (Figure 2D). Similarly, the activity-induced interaction between the enhancer and promoter of *Arc* (Schaukowitch et al., 2014) was also impaired by *Brg1* deletion (Figure 2E).

Cohesin complexes are required for the enhancer-promoter interaction. Our proteomic analysis identified cohesin subunits as BRG1 interacting proteins (Table S1). Using postnatal day 5 (P5) brain nuclear extracts, we confirmed that there is an interaction between endogenous BRG1 and cohesin subunit SMC3 (Figure 2F). By performing cohesin subunit STAG2 ChIP-qPCR, we observed decreased cohesin binding to activity-induced enhancers and promoters in *Brg1^cko^* neurons upon KCl stimulation (Figure 2G). Therefore, BRG1 plays a critical role in modulating enhancer activities by regulating cohesin binding and enhancer-promoter interactions.

It has been shown that neuronal activity induced global accessibility change in adult brain *in vivo* (Su et al., 2017). Previous reports also suggested that BRG1 functions together with the transcription factor AP-1 in enhancer generation during development (Vierbuchen et al., 2017). In cultured neurons, we performed ATAC-seq to identify open chromatin regions and enhancer accessibilities. We identified 9,070 and 5,751 open chromatin sites in basal and depolarized conditions, respectively (Figure S2A). These sites largely overlapped with H3K27Ac identified in neurons (Malik et al., 2014), suggesting that they are active enhancers. Interestingly, neither KCl treatment nor *Brg1* deletion significantly changed ATAC-seq signals (Figure S2B). It is possible that as reported, cultured neurons have a significant different chromatin accessibility state from neurons in vivo (Sinnamon et al., 2019). Thus, in neuron cultures, deletion of *Brg1* impairs enhancer activities but does not significantly change enhancer selection.

### BRG1 phosphorylation at S1382 is induced by neuronal activity

Neuronal activities result in the activation of calcium signaling. The subsequent activation of kinases such as calmodulin-activated CaMKs phosphorylate proteins that mediate neuronal physiological and transcription changes (Deisseroth et al., 2003). To determine how BRG1 and BAF complexes respond to neuronal activities, we affinity-purified endogenous BAF complexes and interactomes from nuclear extracts of resting and depolarized cortical neuron cultures using anti-BRG1/BRM antibody (Figure 3A). Based on silver staining of the purified complexes and mass spectroscopy analyses, there were no obvious subunits differences under the two conditions (Figure 3B and Table S1). Comparison of phosphorylated peptides under basal and depolarized conditions revealed that the peptides corresponding to phosphorylation of BRG1 at S1382 was only detected in the depolarized condition and not in the basal condition (Figure 3C). Phosphorylation of BRG1 at S1382 was demonstrated by proteomic analyses in various human and murine tissues including developing brains (Dephoure et al., 2008; Liao et al., 2008; Mertins et al., 2014). Phosphorylation at this site is dynamic with levels affected by factors including cell-cycle, signaling stimulation, and neural differentiation state. This serine is conserved in *Drosophila* BRM, the only ATPase subunit of the *Drosophila* SWI/SNF complex and a BRG1 homolog, but not in the yeast SWI2/SNF2 protein (Figure 3D). Interestingly, the site in mammalian BRM protein is an alanine despite the close resemblance between BRG1 and BRM in this region of the proteins (Figure 3D). Therefore, the phosphorylation of BRG1 at S1382 may result from a mechanism conserved in *Drosophila* and may be indicative of a function of BRG1 that is distinct from that of BRM.

**Figure 3.**
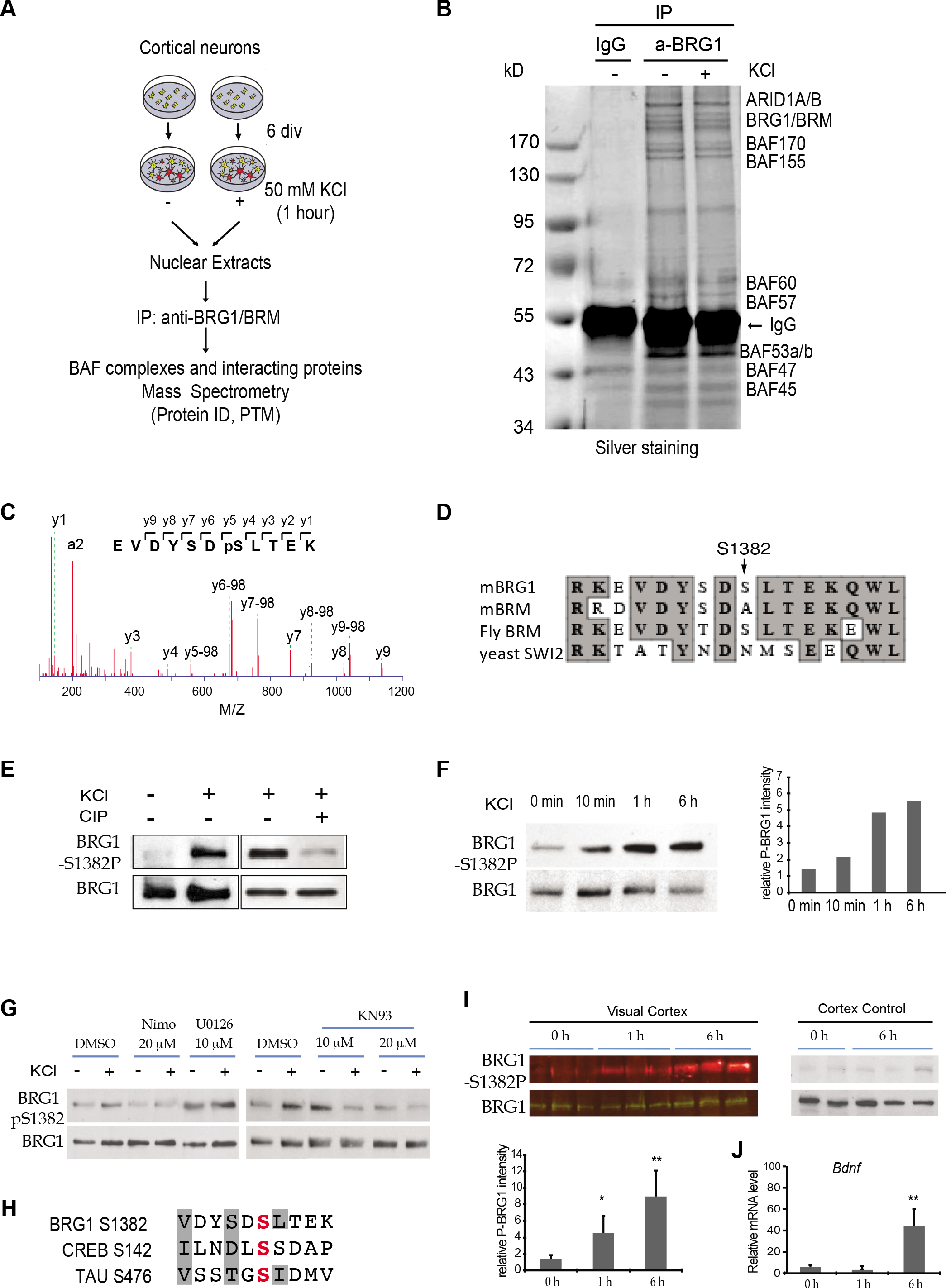
Neuronal activity-induced BRG1 phosphorylation at S1382. A. A schematic of the purification and proteomic analyses of BAF complexes and interacting proteins from cortical neurons under basal and depolarized conditions. B. Photograph of a silver stained gel of affinity-purified neuronal BAF complexes. C. MS/MS fragment ion spectrum with peak assignments for the BRG1 peptide containing phosphorylated S1382. D. Comparison of protein sequences around mammalian BRG1 (mBRG1) S1382 among BRG1 homologs. E. Western blot for KCl-induced BRG1 phosphorylation in cultured neurons and its sensitivity to calf intestinal phosphatase (CIP). F. BRG1 phosphorylation at S1382 in cortical neurons analyzed by western blot after indicated times of KCl treatment. Quantifications are shown on the right. G. Western blot analysis of BRG1 phosphorylation in cortical neurons with and without treatment with indicated inhibitors under basal and depolarizing conditions (KCl 1 h). H. Comparison of protein sequences around BRG1 S1382 and phosphorylation sites (in red) in CREB and TAU. The conserved residues that are important for CamKII are shaded. I. Analysis of light-induced BRG1 phosphorylation at S1382 in mouse visual cortex. Mice were kept in the dark for 3 days and then exposed to light. Western blots show BRG1 phosphorylation before and after light exposure. Cortex controls are anterior one-third of the same cortex. Quantifications are shown below the western blot (n=3 mice per time point). J. RT-qPCR measurement of *Bdnf* mRNA levels in visual cortices after light exposure as described in panel H (n=3 mice per time point). Significance was determined by Student’s t-test. *: p<0.05. **: p<0.01.

We generated an antibody that specifically recognizes S1382-phosphorylated BRG1 in western blot analyses. Using this antibody, we confirmed the increased phosphorylation of BRG1 at S1382 in response to neuron depolarization in culture (Figure 3E). That the observed band corresponds to a phosphorylation event was demonstrated by the decreased in intensity after treatment with calf intestinal phosphatase (CIP) (Figure 3E). Time course studies showed that BRG1 phosphorylation was detected at 10 minutes after KCl stimulation, reached the highest level at the 1-hour time point, and remained high at 6 hours after stimulation (Figure 3F). Using this antibody, we also demonstrated enrichment of S1382-phosphorylated BRG1 in M phase HeLa cells (Figure S3A) in agreement with a previous proteomic study (Dephoure et al., 2008). The antibody also recognized *Drosophila* BRM in its phosphorylated state in S2 cells (Figure S3B).

Neuronal cell depolarization results in Ca^2+^ influx through L-type Ca^2+^ channels, which activates CaMKs and/or MEK/ERK pathways (Deisseroth et al., 2003). We therefore tested the sensitivity of BRG1 phosphorylation to the inhibitors of these pathways. Depolarization-induced BRG1 phosphorylation was sensitive to nimodipine, an L-type calcium channel inhibitor (Cohen and McCarthy, 1987) and to KN93, a CaMKII inhibitor (Sumi et al., 1991), but not to U0126, a MEK/ERK inhibitor (Favata et al., 1998) (Figure 3G). We identified CaMKIIb as a BRG1 interacting protein from our proteomic analyses (Table S1). The region surrounding BRG1 S1382 shares similarities with the phosphorylation sites in several CaMKII substrates such as CREB and Tau (White et al., 1998) (Figure 3H). These results suggest that CaMKII is responsible for BRG1 phosphorylation in response to neuronal activity.

To determine whether BRG1 is phosphorylated in response to physiological stimulation in vivo, we examined BRG1 phosphorylation levels in the mouse visual cortex after light stimulation. At post-natal day 21 mice were put in the dark. After 3 days, mice were exposed to light. At different time points following light stimulation, visual cortices and the anterior part of the cortex control were examined for BRG1 phosphorylation by western blot and for gene activation using RT-qPCR. BRG1 phosphorylation in the visual cortex was significantly increased after light exposure (Figure 3I) and expression of genes such as *Bdnf* was upregulated (Figure 3J). Thus, BRG1 phosphorylation at S1382 is induced by neuronal activation in vivo, possibly through calcium signaling and CaMKII.

### BRG1 phosphorylation is required for regulation of neuronal activity-induced ARG expression

To determine how BRG1 phosphorylation influences ARG expression, we performed rescue experiments with a phosphomimic and a non-phosphorylatable BRG1 mutant at S1382. In KCl-treated *Brg1^cko^* neurons *c-Fos* expression was induced by wild-type and the phosphomimic BRG1-SE. The non-phosphorylatable BRG1-SA failed to rescue impaired *c-Fos* expression (Figure 4A). Expressing BRG1 wild-type or mutant proteins in wild-type neurons had little effects on *c-Fos* expression, possibly due to the difficulty of overexpressing individual BAF subunit in the presence of endogenous BAF complex (Figure 4A). In the *Brg1^cko^* neurons that express BRG1-SA, BRM, which lacks the serine site, or the BRG1 ATPase-inactive mutant BRG1-KR (Khavari et al., 1993), activity-induced expression of *ARGs such as c-Fos and Arc* at 1 hour after KCl treatment was lower than that in cells expressing wild-type BRG1 or BRG1-SE (Figure 4B, 4C). These results indicate that both BRG1 phosphorylation at S1382 and its ATPase activity are required for its function in activating ARGs in neurons. This function is also BRG1 specific and cannot be compensated by BRM. Interestingly, 1 hour after KCl stimulation, in *Brg1^cko^* neurons that express BRG1-SE, LRGs such as *Bdnf* were up-regulated; in contrast, expression of wild-type BRG1 did not result in this up-regulation (Figure 4D). This observation suggests that constitutive phosphorylation of BRG1 S1382 potentiates *Bdnf* enhancers for activation. Therefore, a precise regulation of BRG1 phosphorylation is required to both maintain the repressive basal state and efficient induction of ARGs in response to neuronal signals.

**Figure 4.**
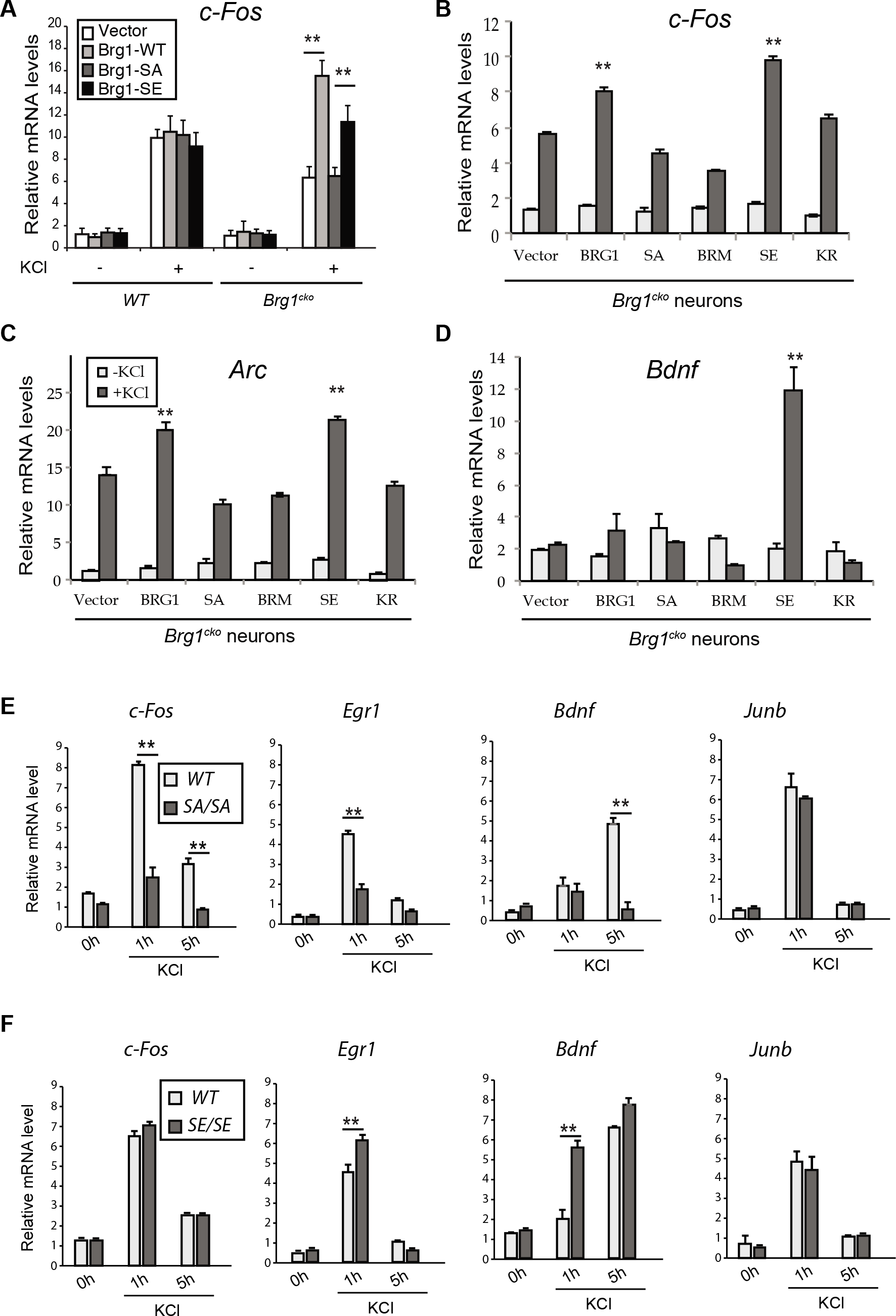
BRG1 S1382 phosphorylation regulates ARG activation. A. RT-qPCR quantification of *c-Fos* mRNA in wild-type and *Brg1^cko^* neurons expressing BRG1-mutant proteins with or without KCl depolarization (1 h). B-D. RT-qPCR analyses of *c-Fos, Arc* and *Bdnf* in *Brg1^cko^* neurons expressing wildtype BRG1 or mutant proteins with or without KCl depolarization (1 h). E. Relative mRNA levels of ARGs in cultured cortical neurons from wild-type and *Brg1-SA* knock-in embryos at different time points after KCl treatment as measured by RT-qPCR. F. Relative mRNA levels of ARGs in cultured cortical neurons from wild-type and *Brg1-SE* knock-in embryos at different time points after KCl treatment as measured by RT-qPCR. Significance was determined by Student’s t-test (n=3). *: p<0.05. **: p<0.01.

### Generating *Brg1-S1382A* and *Brg1-S1382E* knock-in mice

To further understand the function of BRG1 phosphorylation in gene regulation and neuronal function, we generated knock-in mice that harbor the point mutations present in either BRG1-SA or BRG1-SE using the CRISPR/Cas9 method (Figure S4A). The homozygous *Brg1-S1382A* (*SA*) and *Brg1-S1382E* (*SE*) mice express similar levels of BRG1 and BRM and are grossly normal and fertile. Cortical neurons cultured from *SA* mice had impaired activity-induced expression of multiple genes, including *c-Fos, Egr1*, and *Bdnf*, compared to wild-type neurons at 1 and 5 hours after KCl treatment (Figure 4E). *Junb*, which is not a BRG1-dependent IEG, was expressed at the same levels in *SA* and wild-type neurons (Figure 4E, S4B). When we compared wild-type and *SE* cortical neurons, *SE* neurons displayed similar or more sensitive induction of these genes in response to depolarization (Figure 4F, S4C). These results confirm that BRG1 phosphorylation is required for maximum induction of gene activation in response to neuron depolarization and constitutive phosphorylation may sensitize ARGs to induction. We could use these mice to study the biochemical and physiological function of BRG1 S1382 phosphorylation in ARG regulation in vivo.

### BRG1 phosphorylation regulates BAF interactions with transcription co-factors

To understand how BRG1 phosphorylation regulates neuronal gene activation, we examined the biochemical functions of BRG1-SA and BRG1-SE proteins. In SW13 cells where BRG1 and BRM are both absent (Liu et al., 2001), we expressed wild-type and mutant BRG1 proteins using lentiviral vectors. BRG1-SA, BRG1-SE, and BRG1-KR mutants are localized in the nucleus as is the wild-type BRG1 (Figure 5A). In BRG1, S1382 is in the histone-interacting SnAC domain downstream of the ATPase domain (Sen et al., 2013). We affinity-purified BAF complexes containing wild-type and mutant BRG1 and compared their ATPase activities. Unlike the complexes containing ATPase-inactive BRG1-KR, complexes containing BRG1-SA and BRG1-SE had ATPase activities similar to that of the wild-type BAF complex (Figure 5B). Therefore, BRG1 phosphorylation at S1382 does not affect its ATPase activity. Our results described in next section also showed that BRG1 phosphorylation does not affect its binding to chromatin either.

**Figure 5.**
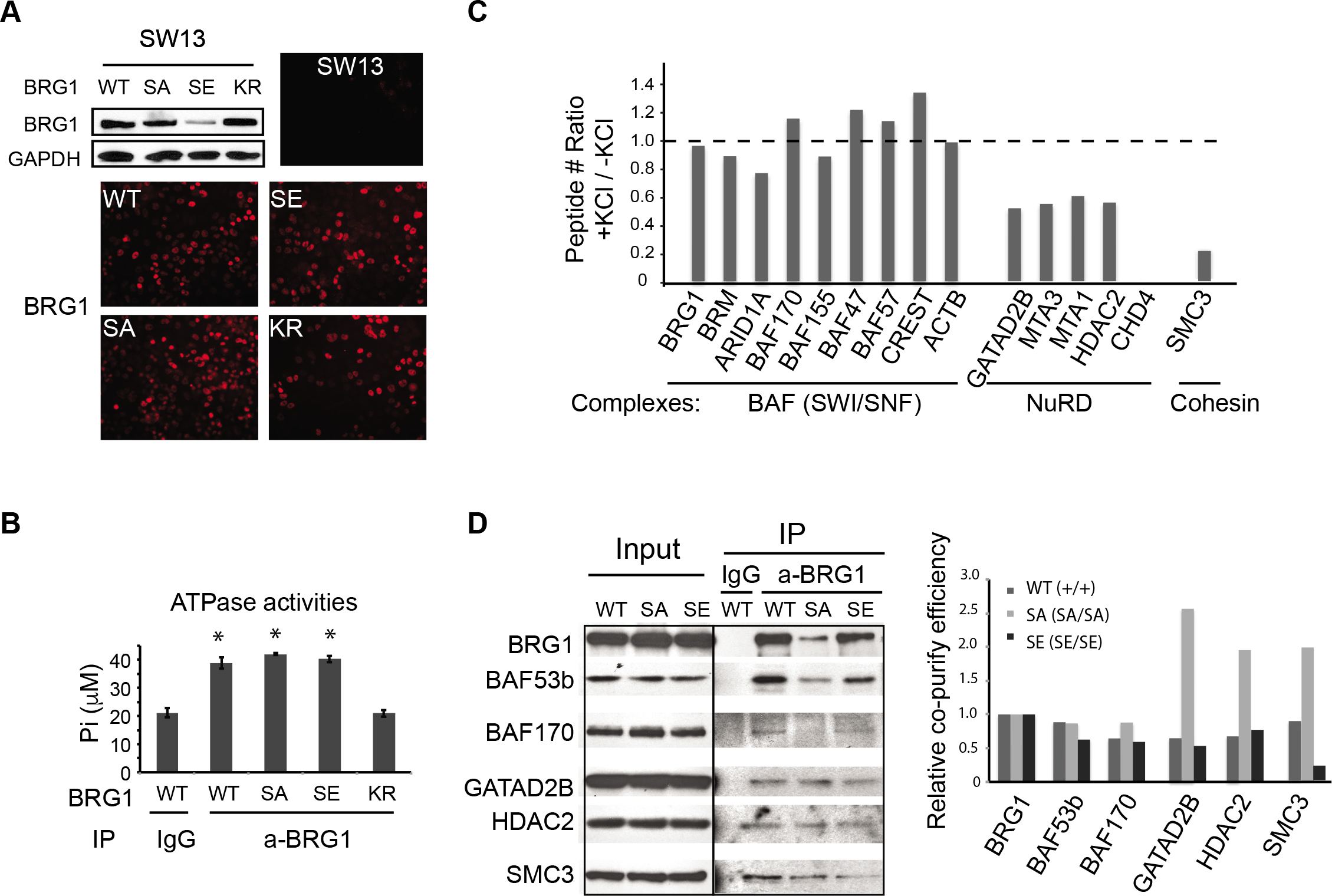
BRG1 phosphorylation affects its interaction with other co-factors without affecting BAF complex formation or ATPase activities. A. Expression and localization of exogenous BRG1 and BRG1-mutant proteins in SW13 cells as detected by western blot and immunostaining. B. ATPase activity assay of BAF complexes purified from SW13 cells expressing BRG1 wild-type or mutant proteins. Significance was determined by ANOVA post hoc t-test (n=3). *: p<0.05. C. Ratios of peptide numbers identified from mass spectrometry for representative BAF subunits, NuRD subunits, and cohesin subunit from proteomic analyses of cortical neurons between depolarized and basal conditions. D. Western blot analyses of samples immunoprecipitated from nuclear extracts from wild-type and *SA* or *SE* P5 cortices using antibodies against BRG1. Quantifications of co-purify efficiency of BRG1 interacting proteins relative to immumoprecipitated BRG1 are shown on the right.

A recent cryo-electron microscopy study of BAF complexes showed that the region of S1382 is likely located at the complex surface (He et al., 2020). To determine whether BRG1 phosphorylation affects its binding to other proteins, we examined the BRG1-interacting proteins identified from our proteomic analyses and compared the abundances of peptides in basal and depolarized conditions. Among the approximately 250 proteins identified, 170 proteins (including all the known BAF subunits and most known interacting proteins) were detected with similar coverage percentages and peptide counts in both conditions (ratios between 0.7 and 1.4; Figure 5C, Table S1). The proteins with peptide number ratios outside of the range may reflect differential interactions with BAF complexes in response to depolarization-induced activation of Ca^2+^ signaling. Notably, four subunits of the NuRD co-repressor complex, GATAD2B, MTA1, MTA3, and HDAC2 (Xue et al., 1998), had relatively high peptide numbers under basal conditions, and all were reduced to 70% or less in the depolarized condition (Figure 5C).

For the NuRD core ATPase subunit CHD4, eight peptides recovered from resting neurons but none from the depolarized neurons (Table S1, Figure 5C). Similarly, four peptides corresponding to cohesin subunit SMC3 were observed in basal conditions but only one was detected in the depolarized condition (Table S1, Figure 5C). Supporting the hypothesis that the interactions between BAF and NuRD and/or cohesin complexes are affected by neuron depolarization and possibly directly by BRG1 phosphorylation at S1382, BRG1 affinity purification from nuclear extracts of cortical tissues from wild-type, *SA*, and *SE* pups at post-natal day 5, when the cortex consists mostly of neurons, showed that although similar ratios of BAF subunits were pulled down from all three genotypes relative to BRG1 levels, more NuRD subunits and SMC3 proteins were copurified with BRG1-SA than wild-type BRG1 or with BRG1-SE (Figure 5D, S5). These results demonstrate that the BRG1 binding affinities for transcriptional co-factors are affected by BRG1 phosphorylation levels. Differences in affinity due to BRG1 phosphorylation may produce distinct ARG transcription outcomes in basal and depolarized conditions.

### BRG1 phosphorylation modulates basal activities of enhancers in the brain

Inducible enhancers are precisely regulated to remain repressed but poised for induction under basal conditions. The basal activities of inducible enhancers may also affect gene induction kinetics in response to signal stimulation. To understand how BRG1 phosphorylation influences basal enhancer activities, we examined levels of H3K27Ac and key transcription regulators at the promoter and the enhancers of *c-Fos* in post-natal day 5 wild-type and *SA* or *SE* cortices. Differences in phosphorylation of BRG1 at S1382 did not affect the amount of BRG1 bound to the *c-Fos* promoter or to enhancer 2 as indicated by BRG1 ChIP (Figure 6A). Interestingly, H3K27Ac ChIP showed that there was a slight but significant increase of H3K27Ac at *c-Fos* enhancer 2 in the cortical samples from *SE* mice compared to wild-type and *SA* mice (Figure 6B), suggesting an increased basal activity of *c-Fos* enhancer in the brain when a BRG1 phosphomimic mutant is expressed than in wild-type cortices or cortices in which BRG1 cannot be phosphorylated. Consistent with this hypothesis, in post-natal day 5 cortex, the cohesin subunit STAG2 was also observed at higher levels on the *c-Fos* promoter and enhancer 2 in *SE* than in *SA* cortices (Figure 6C).

**Figure 6.**
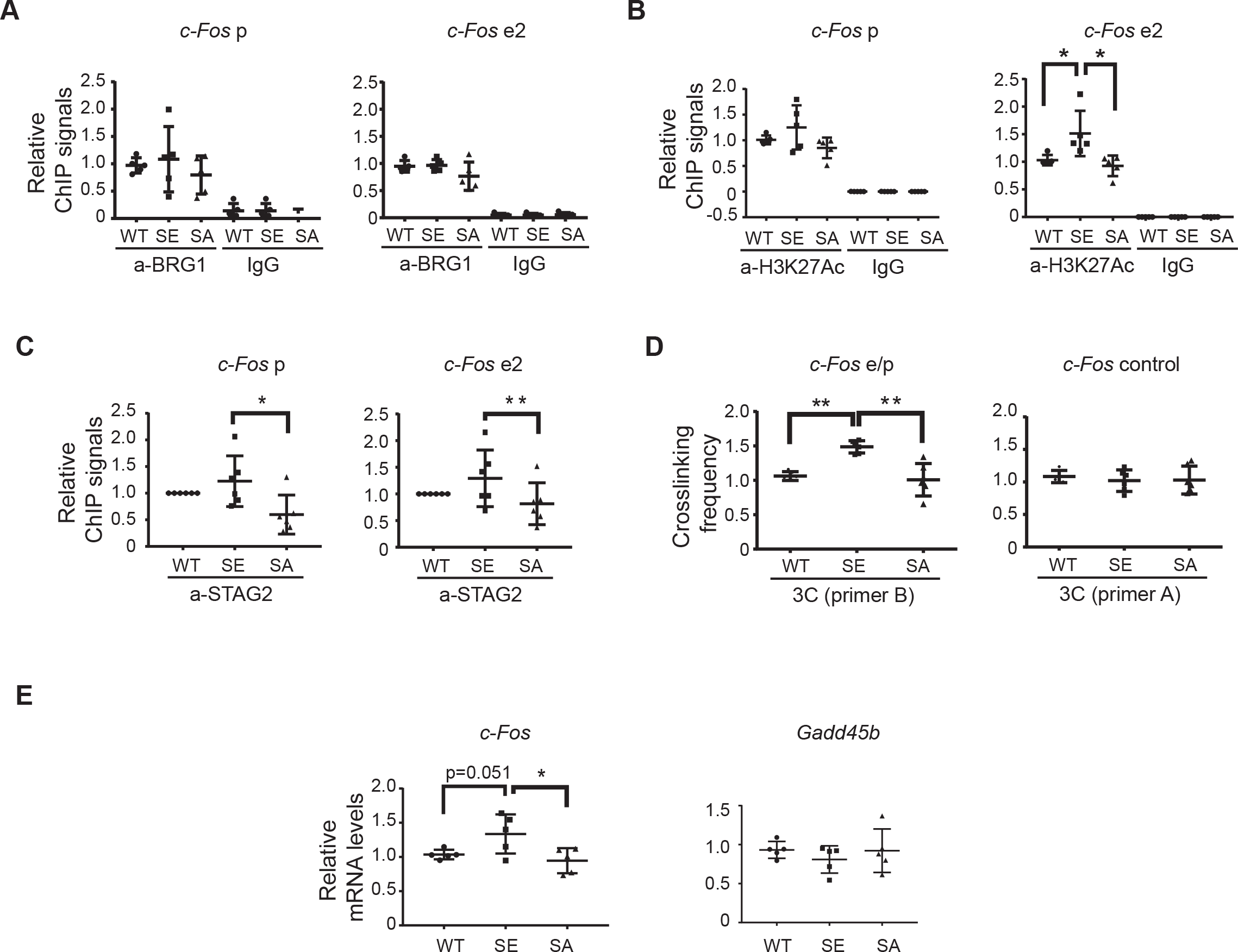
BRG1 phosphorylation regulates *c-Fos* enhancer basal activities in the brain. A. BRG1 and IgG ChIP-qPCR signals at *c-Fos* promoter (left) and enhancer (right) regions in P5 *SA* and *SE* cortices relative to signals in wild-type tissue. B. H3K27Ac and IgG ChIP-qPCR signals at *c-Fos* promoter (left) and enhancer (right) regions in P5 *SA* and *SE* cortices relative to signals in wild-type tissue. C. STAG2 and IgG ChIP-qPCR signals at *c-Fos* promoter (left) and enhancer (right) regions in P5 *SA* and *SE* cortices relative to signals in wild-type tissue. D. Crosslinking frequency between *c-Fos* enhancer and promoter (left) and control regions (right) in P5 *SA* and *SE* cortices measured in a 3C experiment relative to signals in wild-type tissue. E. Levels of *c-Fos* and control *Gadd45b* mRNAs in P5 wild-type and *SA* and *SE* cortices measured by RT-qPCR. Significance was determined by ANOVA post hoc t-test (n=5 pups per genotype). *: p<0.05. **: p<0.01.

We next performed 3C experiments to examine the interaction between *c-Fos* enhancer 2 and the *c-Fos* promoter. A significantly higher interaction rate was observed in *SE* cortical samples than in *SA* and wild-type samples (Figure 6D). Collectively, these results indicate that there is a higher basal *c-Fos* enhancer activity in *SE* mutant neurons. The higher enhancer activities caused a slight but significant increase of basal expression of *c-Fos* in *SE* cortices than wild-type or *SA* cortices as shown by RT-qPCR (Figure 6E). Therefore, an unphosphorylated BRG1 is required to repress basal *c-Fos* enhancer activities and BRG1 phosphorylation facilitates the de-repression and efficient activation of this ARG in response to neuronal activities. Although BRG1 phosphorylation alone is not sufficient to induce high levels of gene activation, it increases the basal activities of inducible enhancers, which may facilitate gene activation in response to signal stimulation.

### BRG1 phosphorylation regulates brain IEG expression in response to stress and mutants display anxiety-like behaviors

ARG expression affects many aspects of neuronal function and behaviors, especially in response to stress (Gallo et al., 2018). To determine whether BRG1 phosphorylation is important for in vivo neurological functions, we performed behavioral tests of the *SA* and *SE* mice. Both *SA* and *SE* mice were generated and maintained in the C57Bl/J6 background. In the light-dark box test, both *SA* and *SE* mice tended to stay for less time in the lit region than did wild-type C57Bl/6J mice (Figure 7A). *SE* mice were also significantly less active than wild-type mice in the light box, but displayed normal activities in general (Figure 7A). In elevated maze test, both *SA* and *SE* mice spent less time in open arms than did wild-type mice (Figure 7B). Further, *SA* mice entered the open arms less frequently and spent significantly more time in the closed arm than wildtype and *SE* mice (Figure 7B). These results indicate that alterations in BRG1 phosphorylation cause anxiety-like behaviors. These defects could be caused directly by altered expression of ARGs or indirectly by altered neural functions due to BRG1 defects during development.

**Figure 7.**
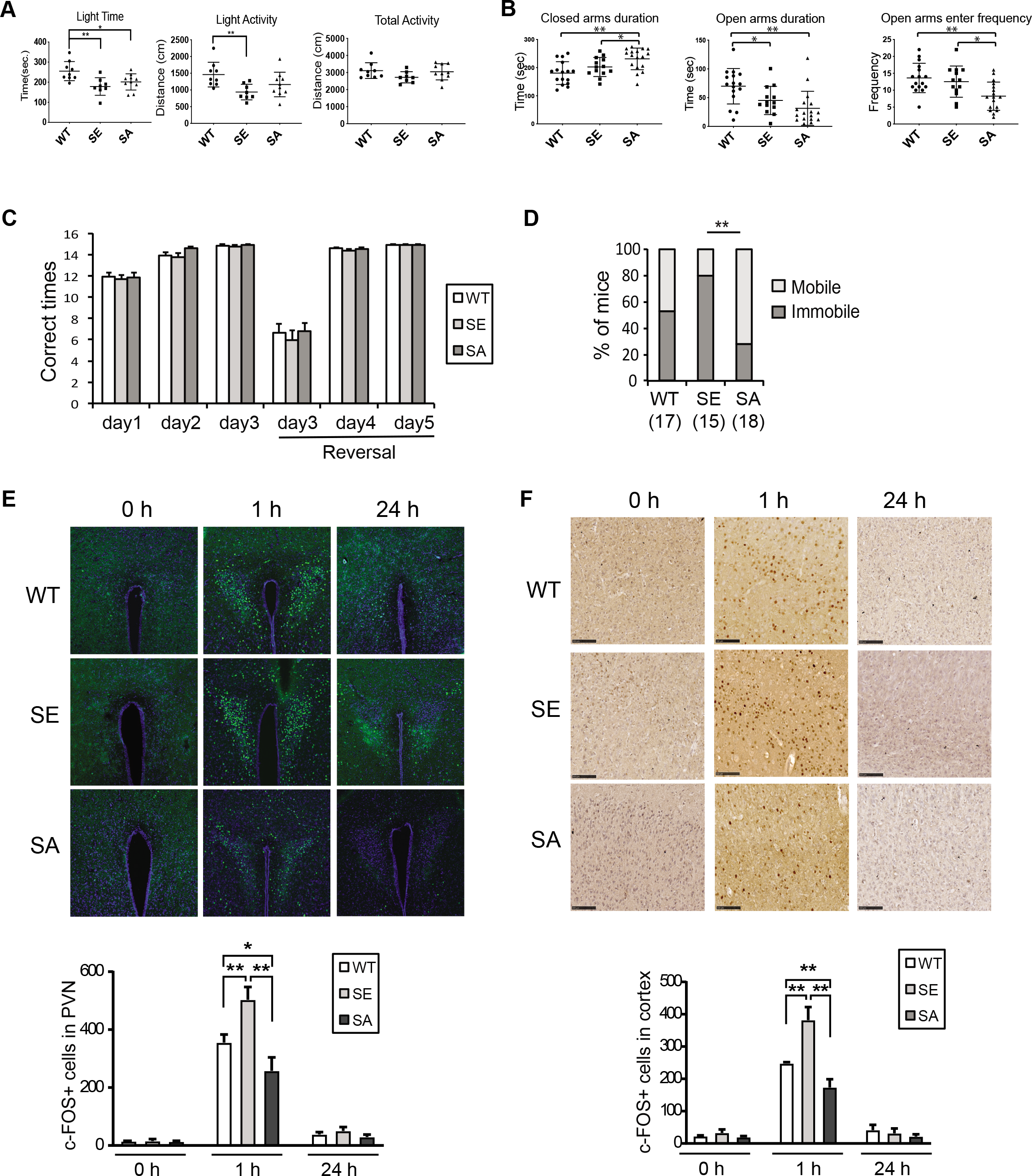
Alteration of BRG1 phosphorylation states causes anxiety-like behaviors and alterations in *c-Fos* expression in response to stress. A. Time spent in the light box in the light-dark box test, distance traveled in the light box and total distance traveled for wild-type (n=17), *SA* (n=15), and *SE* (n=18) mice. B. Time spent in open and closed arms of the elevated maze and open arm entering frequency for wild-type (n=17), *SA* (n=15), and *SE* (n=18) mice. C. Number of times out of 15 trials that wild-type (n=17), *SA* (n=15), and *SE* (n=18) mice found the platform in each session of the Y maze reversal learning swimming test. D. The percentage of mice that were immobile for longer than 30 seconds once put in water in any trials for wild-type (n=17), *SA* (n=15), and *SE* (n=18) groups. Two proportion Z-test, **: p<0.01. E. Representative images of C-FOS-stained paraventricular nucleus (PVN) regions of brain sections of mice before and after swim test. Quantification of C-FOS-positive cells in PVN is shown below the images (n=3). F. Representative images of C-FOS-stained somatosensory cortex regions of brain sections of mice before and after swim test. Quantification of C-FOS-positive cells is shown below the images (n=3). Significance (A, B, E and F) was determined by ANOVA post hoc t-test. *: p<0.05. **: p<0.01.

In a Y maze reversal learning swimming test, both *SA* and *SE* mice learned to find the correct targets in the initial and reversal learning phases with frequencies similar to that of wild-type mice (Figure 7C). Interestingly in the first several trials on the first day of swimming training, all mice started swimming shortly after being put in water. However, after several rounds of swimming over a time period of about 1-2 hour, some mice became immobile for longer than 30 seconds in subsequent trials, possibly due to stress responses. When the mice that displayed immobile behaviors were counted, there is a significant difference among different genotypes. *SE* mice showed highest percentage with this phenotype, whereas *SA* mice showed lowest percentage (Figure 7D). We examined mouse brains for c-FOS expression in various regions important for stress response at 1 hour after the swimming test. We observed consistently higher c-FOS expression in *SE* brains in regions such as the hypothalamus paraventricular nucleus and somatosensory cortex than were observed in wild-type brains, whereas *SA* brains expresses less c-FOS than wildtype (Figure 7E, 7F). Since over expression of IEGs have been linked to anxiety and altered stress responses (Gallo et al., 2018; Yap and Greenberg, 2018), it is likely that BRG1 phosphorylation states regulate these neurological behaviors through a direct impact on ARG expression. The elevated basal *c-Fos* enhancer activities in *SE* brains would result in the stronger induction of *c-Fos* expression, whereas *c-Fos* induction is impaired in *SA* brains without BRG1 phosphorylation. Thus, BRG1 phosphorylation is important for regulation of BRG1-mediated enhancer activation and ARG induction during neuronal responses to stress and stimulation.

## DISCUSSION

In this study, we uncovered molecular mechanisms underlying the function of SWI/SNF chromatin remodeler BRG1 in enhancer activation in response to neuronal activity (Figure S6). We identified an activity-induced phosphorylation event that modulates BRG1 function in enhancer regulation. We demonstrated that unphosphorylated BRG1 is required to maintain enhancers in a repressed state under basal conditions and that BRG1 phosphorylation in response to neuronal activities facilitates the activation of these signaling-inducible enhancers. BRG1 phosphorylation affects its interactions with important transcription co-factors, and we observed defects in stress-induced ARG expression and anxiety-like behaviors in mice that expressed constitutively phosphorylated or non-phosphorylatable BRG1. Thus, we revealed a novel level of regulation of BRG1 responsible for fine-tuning of signal-induced enhancer activation and gene expression.

BRG1 and BAF chromatin remodeling complexes are enriched at enhancer sites during development and in certain disease states (Alexander et al., 2015; Alver et al., 2017; Hodges et al., 2018). BAF complexes have context-dependent functions and can act as activators or as repressors depending on the subunit composition, chromatin environment, and interacting epigenetic regulators (Hodges et al., 2016; Kadoch, 2019; Wu, 2012; Wu et al., 2009). In our study, we took advantage of the relatively synchronized ARG induction in response to KCl-induced depolarization to evaluate BRG1 function in enhancer activation. A previous study showed that RNAi-mediated inhibition of BRG1 expression in cultured cortical neurons resulted in the induction of *c-Fos* expression at a lower concentration of KCl than observed when normal levels of BRG1 were present (Qiu and Ghosh, 2008b). Our observations that BRG1 represses basal activities of ARG enhancers are consistent with this (Figure 6). In addition, we found that BRG1 is required for signal-induced ARG activation. Our previous study also showed that BRG1 plays a dual role in sonic hedgehog-mediated signaling by repressing gene expression under basal conditions and activating signaling-induced gene expression (Zhan et al., 2011). Therefore, BRG1-containing BAF complexes could set up the precise basal activity and inducibility necessary for the proper function of inducible enhancers. Interacting protein change mediated by signal-induced phosphorylation could be one mechanism that modulates BRG1 function.

Recruitment of BAF complexes to enhancers is thought to be mediated through interactions with transcription factors that bind sequence-specifically to enhancer regions. We have previously shown that MEF2C is required for activity-induced BRG1 binding to the promoters of several ARGs (Zhang et al., 2016). It has been also shown that AP-1 interacts with BRG1 and is possibly important for enhancer determination (Vierbuchen et al., 2017). In addition to transcription factors, the histone acetyl transferase p300/CBP may recruit BAF to enhancers (Alver et al., 2017). We showed that BRG1 binding to enhancers is sensitive to inhibitors of its bromodomain and to inhibitors of p300/CBP (Figure 1G-I). This result suggests that an interaction between the BRG1 bromodomain and H3K27Ac contributes to activity-induced recruitment of BRG1 to enhancers. Thus, a specific and precise targeting of BAF complexes to enhancers is likely dependent on the coordinated work of BAF interacting transcription factors, co-factors and chromatin features.

Besides transcription factors, transcription co-factors such as the cohesin complex that facilitate enhancer-promoter looping, histone modification enzymes such as p300/CBP and HDACs, chromatin regulators such as BAF and repressor complex NuRD, mediator complexes, RNA polymerase II, and enhancer RNAs all contribute to enhancer activities and ultimately mRNA expression. However, how these co-factors coordinate with each other to produce precise transcriptional outcomes is not clear. For example, super enhancers could organize the formation of large protein complexes held together by multiple low-affinity interactions between low-complexity regions in transcription factors and co-factors to induce high levels of gene expression (Sabari et al., 2018). The balance and competition between antagonizing co-factors also may underlie the precise regulation of the enhancer activities and gene expression. We discovered that BRG1 plays a central role in coordinating interactions of transcription co-factors at enhancers to maintain precise activities under basal and activated conditions. *Brg1* deletion in activated neurons reduced cohesin binding, enhancer-promoter looping, H3K27Ac levels, RNA polymerase II recruitment, and eRNA expression levels for the *c-Fos* and *Arc* genes, two representative ARGs (Figure 2). BRG1 recruitment also relies on the rapid increase of H3K27Ac, and it has been shown that eRNA stimulates p300/CBP HAT activities (Bose et al., 2017). Therefore, although at certain transcription activation stages, the recruitment of different co-factors could be sequential, the enhancer activation process depends on the coordinated action of transcription factors and cofactors, and BRG1 is an essential player.

We identified a dynamic BRG1 phosphorylation site at S1382 that affects BRG1 interactions with several transcription co-factors. We showed that the interaction of BRG1 with the NuRD complex was reduced by phosphorylation of BRG1 at S1382 (Figure 5C, 5D). The NuRD complex is an important co-repressor that has been shown to bind to enhancers and to antagonize BAF complexes to fine-tune enhancer activities (Bornelov et al., 2018; Bracken et al., 2019; Gao et al., 2009). In cerebellar granule neurons, NuRD complexes regulate ARG expression after activation (Yamada et al., 2014; Yang et al., 2016). Therefore, phosphorylation-mediated regulation of the dynamic interactions of BRG1 with NuRD complexes could underlie the dual functions of BRG1 in repressing basal levels and activating signaling-induced ARGs (Figure S6). Another important enhancer regulator that displayed differential interaction with BRG1 upon phosphorylation is cohesin subunit SMC3. Interestingly, although *Brg1* deletion led to decreased activity-induced cohesin binding to enhancers, the interaction of BRG1 with cohesin was reduced by neuronal activation and by BRG1 phosphorylation at S1382 (Figure 5C, 5D). Therefore, it is not likely that BRG1 recruits cohesin to enhancers. Cohesin dynamics have been shown to be important for its function in mediating enhancer-promoter interactions (Haarhuis et al., 2017; Kueng et al., 2006), and the reduced interactions with BRG1 upon depolarization may increase cohesin dynamics. Further studies will be needed to investigate how BRG1 phosphorylation affects the activities of these enhancer regulators.

BRG1 has been shown to be phosphorylated during mitosis, which correlates with its dissociation from condensed chromatin and inactivation (Muchardt et al., 1996; Sif et al., 1998). Recent studies showed that Casein kinase 2 phosphorylates BRG1 during mitosis at multiple sites, and that these phosphorylation events are important for myoblast proliferation and survival (Padilla-Benavides et al., 2020; Padilla-Benavides et al., 2017). During myogenesis, phosphorylation of BRG1 by PKCβ1 and dephosphorylation by calcineurin regulates gene expression (Nasipak et al., 2015). In *Drosophila*, phosphorylation of BRM by cyclin-dependent kinases is necessary for cell proliferation and development of wing epithelium (Roesley et al., 2018). However, it remains unclear how these phosphorylation events regulate transcriptional activities of BRG1 and BRM at the molecular level. In addition to BRG1 phosphorylation during cellcycle progression and development, BRG1 has been shown to be phosphorylated upon DNA-damage signaling by the kinase ATM, resulting in the binding of the ATPase to γ-H2AX containing nucleosomes (Kwon et al., 2015). Phosphorylation of BRG1 at S1382 has been characterized in multiple unbiased proteomic analyses in rodent and human cells (Dephoure et al., 2008; Liao et al., 2008; Mertins et al., 2014). The phosphorylated form of BRG1 is enriched in M phase and is also observed in tumor cells and in the developing brain. We found that BRG1 phosphorylation at S1382 in postmitotic neurons is regulated by calcium signaling pathways mediated by CaMKII. We also provided biochemical and genetic evidence supporting a function of BRG1 phosphorylation in regulating the interaction with other co-factors and in regulating enhancer activities. In addition to CaMKII in neurons, it is possible that other kinases and phosphatases could regulate BRG1 phosphorylation to modulate enhancer activities in response to various signaling and cell cycle progression during normal development and in cancers.

Finally, using knock-in mice we demonstrated that phosphomimic and non-phosphorylatable BRG1 mutations both led to abnormal neuronal responses to stress, changes in IEG expression, and anxiety-like behavior, indicating the importance of BRG1 phosphorylation for normal neuronal development and functions in vivo. Mice with phosphomimic and non-phosphorylatable BRG1 mutations are grossly normal, indicating that BRG1 phosphorylation is not required for overall brain development. The behavioral defects could be caused by alterations in IEG or ARG expression in neurons or could be due to altered neural circuits or abnormal neuronal activities that result from impaired ARG expression during development. It has been shown that neuronal activities are important for the development of visual system (Berry and Nedivi, 2016; Wiesel, 1982) and that ARG expression is important for synaptic plasticity (West and Greenberg, 2011; Yap and Greenberg, 2018). It would be interesting to further characterize the mouse strains with mutations in BRG1 to determine the short-term and long-term effects of ARG expression changes.

In summary, this study not only demonstrated that BRG1 functions to coordinate chromatin regulation of enhancer activities but also identified a signaling pathway that regulates ARG expression through BRG1 phosphorylation. These functions are important for normal neural responses to stress. Both *Brg1* mutations and altered BRG1 phosphorylation may contribute to human neurological and psychiatric diseases.

## Acknowledgements

We thank Dr. Taekyung Kim for providing 3C reagents, Dr. Hongtao Yu for providing the anti-STAG2 antibody, Dr. Shari Birnbaum and the UTSW Rodent Behavior Facility for aiding in the behavior tests, and the UTSW Sequencing Facility for performing the nextgeneration sequencing. We thank Mr. Huaxia Dong for technical help with mouse breeding. This work was supported by grants from the Welch Foundation (I-1940-20170325 to J.W.) and the NIH (R01NS09606 and R21NS104596 to J.W.).

## Contributions

J.W., B.K., and Y.L. designed the experiments. B.K., Y.L., X.Z., Z.Z., X.S., and J.Y. performed the experiments and collected the data. B.K., Y.L., X.Z., Z.Z., and J.W. analyzed the results. Z.X. performed the main bioinformatics analyses. J.W. wrote the manuscript with help from all authors.

## METHODS

### Mice

*Brg1^F/F^ BAF53b-Cre (Brg1^cko^*) and control embryos (Zhang et al., 2016) were generated from crosses between *Brg1^F/+^ BAF53b-Cre* and *Brg1^F/F^* mice and were maintained on a mixed genetic background. *Brg1-SA (SA*) and *Brg1-SE (SE*) mice were generated by injecting Cas9 recombinant protein, gRNA targeting the 5’-ccaCC**GC**AAGGAGGTAGACTACA-3’ site upstream of the codon encoding S1382 in BRG1, and repair single-stranded DNA oligonucleotides into C57BL/6 pronuclei. Progenies were screened by sequencing the target sites (Figure S4A) and back-crossed to wild-type C57BL/6 mice to obtain heterozygous mice and subsequent homozygotes. All mice are housed at UT Southwestern Medical Center Animal Facility. All procedures were performed in accordance with the IACUC-approved protocols. In all animal experiments, both males and females were used, and there were no significant differences found between genders.

### Plasmid construction, virus preparation, and transfection or infection

The constructs for expression of BRG1, BRM, BRG1-SA, BRG1-SE, and BRG1-KR were generated by inserting the coding regions of these genes into pSin4-EF2-IRES-Puro lentiviral vectors (Zhan et al., 2011). Lentiviruses were propagated in HEK293T cells according to a previously described procedure (Zhan et al., 2011), followed by ultracentrifugation for concentration. Attached cultured SW13 and primary neurons were infected at an MOI of 5 for 24-48 h. PolyJet (Signagen) was used for plasmid transfection of cultured cells.

### Cortical neuron culture and inhibitor treatment

E16.5 cortical cells were cultured as previously described (Wu et al., 2007; Zhang et al., 2016). Dissociated cortical neurons were plated on poly-L-ornithine- and fibronectin-coated wells. Culture media contained neurobasal plus with B27 plus supplement (Gibco). Cortical cultures were infected with lentiviruses on 4 days *in vitro* (div) and analyzed at 6 div. For depolarization, the cultures were pretreated with TTX (1 μM) and APV (100 μM) to inhibit spontaneous activation on 5 div and 50 mM KCl was added to the cultures for 1 to 6 h on 6 div as described (Flavell et al., 2006). For inhibitor treatment, PFI-3 (final concentration 20 μM), C646 (25 μM), or DMSO (as a control) was added 4 h before KCl treatment on 6 div. Nimodipin, KN93, and U0126 treatments were performed with indicated concentrations on 5 div before KCl treatment on 6 div.

### RT-PCR and q-PCR

RNAs from cells or ground tissues were extracted with TRIZOL (Invitrogen). cDNAs were synthesized by reverse transcription using iScript (Bio-Rad), followed by PCR or quantitative PCR analysis. A Bio-Rad real-time PCR system (C1000 Thermal Cycler) was used for quantitative PCR. Levels of RNAs of interest were normalized to *GAPDH* mRNA. Bar graphs shown are representative of experiments performed in triplicate unless otherwise indicated. The experiments were repeated at least three times. Dot graphs represent results from individual samples. Standard errors were calculated according to a previously described method (Zhan et al., 2011). The sequences of all the primers are listed in Table S2.

### ChIP experiments and ChIP-seq analyses

ChIP experiments were performed as described previously (Shi et al., 2016; Zhan et al., 2011). Dounced tissue or dissociated cells were crosslinked with PFA or double crosslinked with DSG (Pierce), and nucleic acids were sonicated into fragments of 200-500 bp. Antibodies used were against BRG1/BRM (J1) (Khavari et al., 1993), H3K27Ac (Abcam, ab4729), RNA pol II (Abcam, ab817), and STAG2 (a gift from Dr. Hongtao Yu, UTSW). J1 antibody has been used previously for BRG1 ChIP-seq analyses (Ho et al., 2009; Yu et al., 2013). Precipitated DNA was purified and subjected to either real-time PCR (Table S2) or next generation sequencing. NEBNext ChIP-Seq Sample Prep Master Mix Set 1 was used for library generation, and a Hiseq 2500 sequencer was used for sequencing at the UT Southwestern Medical Center Sequencing Core Facility. Short reads were mapped to UCSC reference mouse genome (GRCm38/mm10) with BWA aligner (Li and Durbin, 2009) and then SICER was used to detect the BRG1-binding regions (Shi et al., 2016; Zang et al., 2009). Default parameter settings with three 200-bp windows were used to calculate the enrichment of BRG1-binding regions. The corresponding input sample was used as control. Duplicate reads were removed before peak calling by SICER. Statistically significant peaks (FDR<0.05) enriched in the BRG1-ChIP sample relative to its corresponding input sample were annotated for genomic location. H3K27Ac ChIP-seq data was downloaded from NCBI GEO (GSE60192) (Malik et al., 2014), and aligned with UCSC reference mouse genome (GRCm38, Dec. 2011) with BWA, following peak calling by MACS2 (Zhang et al., 2008) with default parameter to call narrow peaks. DeepTools (Ramirez et al., 2016) was used to illustrate the reads distribution from ChIP-seq in the enhancers and their 10-kb flanking regions. BRG1 ChIP and H3K27Ac ChIP peaks were overlapping if there was 1 bp overlap of the peak regions.

### Immunoprecipitation and western blot

Cortical tissues or cultured cortical cells were harvested and lysed in buffer A (25 mM Tris, pH 7.5, 25 mM KCl, 5 mM MgCl, 0.1% NP-40, 10% glycerol). Nuclear pellets were resuspended in RIPA buffer (25 mM Tris, pH 7.5, 150 mM NaCl, 0.05% SDS, 1% Triton X-100, 0.5% SDC) to prepare nuclear extracts. The anti-BRG1 J1 antibody was preincubated with Protein A-coated magnetic Dynabeads (Invitrogen) before adding to nuclear extracts. Samples were incubated at 4 °C overnight, beads were washed with RIPA buffer four times. Precipitated proteins were eluted by boiling in 2X Sample Buffer (Bio-Rad) before SDS-PAGE and western blot analysis. Protease and phosphatase inhibitors were used in all immunoprecipitation buffers. For immunoblotting, cell lysates or precipitated proteins were separated on SDS-PAGE gels. Antibodies used were against BRG1 (G7, Santa Cruz Biotechnology), BRG1/BRM (J1), SMC3 (E3, Santa Cruz Biotechnology), GAPDH (G9545, Sigma), BAF53b (Wu et al., 2007), BAF170 (Wang et al., 1996), GATAD2B (NB100-60646,Novus Biologicals), HDAC2 (PA-1-861, Invitrogen), H3S10P (#9701, Cell Signaling). HRP-conjugated (Jackson Immunology) or IRDye-conjugated secondary antibodies (LI-COR) were used in western blot. Rabbit BRG1-S1382 phospho-specific antibody was generated against peptide antigen VDYSD(pS)LTEKQ by Ab-Mart (Shanghai).

### ATPase activity assay

ATPase activity was measured using a high-sensitivity colorimetric ATPase assay kit following the manufacturer’s instruction (Innova Biosciences). Briefly, affinity-purified BAF complexes as described above were suspended in 25 mM Tris, pH 7.5 and incubated with the reaction buffer (100 mM Tris, pH 7.4, 5 mM MgCl_2_, and 1 mM ATP) in the presence of 0.5 μg plasmid DNA for 15 min at 37 °C. The reaction was stopped by adding PiColorLock mix. The amount of inorganic phosphate released was quantified colorimetrically at 620 nm.

### Affinity purification of BAF complexes and mass spectrometry analysis

Nuclear extracts are prepared as describe above from 6-div mouse cortical neurons with or without KCl treatment for 1 h. J1 antibody or rabbit IgG control was cross-linked to GE FastFlow Sepharose Protein A beads as previously described (Lessard et al., 2007). BAF complexes were affinity purified from nuclear extracts (600 μg) using 20 μg antibody-conjugated beads. Protein complexes were eluted with 0.1 M acetic acid. After silver staining (LC6070, Invitrogen) to confirm purity, the eluted BAF complexes and associated proteins from gel slices were subjected to LC-MS/MS, which was performed by PTM Biolabs. Protein mixes were digested in the gel with trypsin, and peptides were extracted. The peptides were separated and then analyzed using a Q Exactive Plus Hybrid Quadrupole-Orbitrap mass spectrometer (ThermoFisher Scientific). The resulting MS/MS data were processed using the Mascot search engine (version 2.3). Tandem mass spectra were searched against SwissProt database concatenated with reverse decoy database. Trypsin/P was specified as cleavage enzyme allowing up to 3 missing cleavages. Mass error was set to 10 ppm for precursor ions and 0.02 Da for fragment ions. Phosphorylations of Ser, Thr, and Tyr were specified as variable modifications. An FDR of less than 1% was used to filter the identified peptides. All the other parameters in Mascot were set to default values. We identified 272 proteins with 35 phosphorylation sites in the basal condition and 305 proteins with 40 phosphorylation sites in depolarized neuron samples.

### Chromosome conformation capture

Chromosome conformation capture (3C) experiments were performed as described previously (Naumova et al., 2012). Extracts of cultured neurons or homogenized mouse cortices collected at post-natal day 5 (P5) were crosslinked with 1% formaldehyde for 10 min; the reaction was stopped by addition of 0.125 M glycine. Crosslinked cells were lysed with in buffer (10 mM Tris, pH 8.0, 10 mM NaCl, 0.2% NP-40) containing protease inhibitor for 15 min at 4 °C. Nuclei were resuspended in the 1X NEB restriction enzyme buffer, and 1% SDS was added. After incubating for 10 min at 65 °C, 1% Triton X-100 was added. Samples were digested with SacI for *c-Fos* analysis or with BgIII and NcoI for *Arc* analysis at 37 °C overnight. The restriction enzyme was inactivated with 1.6% SDS for 30 min at 65 °C followed by incubation in 1% Triton X-100 for 1 h at 37 °C. Ligation was performed using T4 DNA ligase for 4 h at 16 °C and then 30 min at room temperature. Samples were treated with 200 μg proteinase K at 65 °C overnight for reverse crosslinking, followed by RNase treatment (1 μg/ml) for 1 h at 37 °C. After phenol-chloroform extraction and ethanol precipitation, DNA was analyzed by qPCR using primers described previously (Joo et al., 2016) (Table S2). Two bacterial artificial chromosome clones containing the *c-Fos* and *Arc* genes were used as negative controls (Joo et al., 2016).

### ATAC-sequencing

Cultured wild-type and *Brg1^cko^* cortical neurons with or without KCl treatment for 1 h were harvested in PBS. Neurons (50,000 cells/sample, n=2 in each condition) were lysed with lysis buffer (10 mM Tris, pH 7.4, 10 mM NaCl, 3 mM MgCl_2_, 0.1% NP-40). The transposition reaction was performed using Tn5 transposase from the Nextra kit (Illumina, FC-121-1030) at 37 °C for 30 min. Transposed DNA samples were purified using a Qiagen MinElute Kit (catalog no. 28004) followed by PCR amplification. Sequencing libraries were generated using Ad1_noMX with barcoded primers and were amplified (7-9 cycles) before DNA purification and concentration. All samples were sequenced using a Hiseq 2500 sequencer with 75-bp paired end reads and 60 million reads were obtained for each library. The in-house script was developed to remove the adapter and extract the genomics sequences from raw sequencing reads. Bowtie 2 (Langmead and Salzberg, 2012) was used to align all reads to UCSC reference mouse genome (GRCm38/mm10), and open-chromatin regions were called by MACS2 (Zhang et al., 2008) with parameter of “callpeak --nomodel --shift −100 --extsize 200.”

### Behavior tests

All experiments in this study were performed in the Behavior Core Facility and approved by the IACUC at UT Southwestern Medical Center. Mice were housed with food and water available *ad libitum* with a 12-h light/dark cycle, and all behavior testing occurring during the light cycle. Behavioral tests were conducted by testing less stressful behaviors before more stressful ones.

Light/Dark box test: Mice were placed in the dark chamber of a custom-made light/dark box and allowed to habituate for 2 min. After opening of the divider separating the dark side from the light side, mice were allowed to freely explore both chambers for 10 min while monitored from above by a video camera connected to a computer running video tracking software (Ethovision 3.0, Noldus) to measure the time, frequency, and activity (travel distance) in each chamber. Each chamber of the box was 25 × 26 cm with 1,700 lux on the light side and ~0.1 lux on the dark side.

Elevated maze test: Mice were placed in the center of a white Plexiglas elevated plus maze (each arm 33 cm long and 5 cm wide with 15-cm-high black Plexiglas walls on closed arms) and allowed to explore for 5 min. The test was conducted in dim white light (~7 lux). Data on duration of time spent in each arm and entering frequencies were recorded and analyzed using the CleverSys TopScan software.

Y maze reversal learning: The Y maze (each arm 7 cm wide and 20 cm long) was filled with water (approximately 24 °C) and a small amount of non-toxic paint. One arm of the maze was designated the Start Arm, the other two arms are the Choice Arms. One of the Choice Arms had a platform located at the end, submerged 1-2 cm below the water level. Mice were placed into the Start Arm of the maze and allowed to swim to locate the hidden platform. Once the mouse found the platform, it was removed from the maze. Mice that did not locate the platform within 60 sec were gently guided to the platform and then removed from the maze. Mice were placed briefly on a paper towel to dry after removal from the maze. Each mouse was placed in the maze 15 times per day with at 1-30 min between trials. The mice were trained for 3 days. On the next day, the platform location was moved to the opposite arm, and each mouse was tested an additional 15 times. The number of times the mouse found the platform each day was recorded. During the experiment, certain mice remained immobile in the water unless stimulated. We defined immobile mice as those that floated but did not swim for longer than 30 s after being put in water for any trial. Immuno-fluorescent staining of c-FOS was performed using c-FOS antibody (9F6, Cell Signaling) on brain cryosections of mice that had performed six trials in 1 h. c-FOS-positive cells were counted from paraventricular nucleus and cortical regions from three mice (3 sections/brain), and averages are reported.

### Statistical analysis

Unless otherwise specified, at least three independent experiments were performed and each condition was analyzed in triplicate. Bar graphs shown are representative experiments. Dot graphs represent results from individual samples. Data are expressed as means ± s.d. Statistical analysis was performed by either analysis of variance with ANOVA post hoc t-test for multiple comparisons or a two-tailed unpaired Student’s t-test. A p value of <0.05 was considered significant.

