## Supplemental figures and tables for "Neuronal Activity-Induced BRG1 Phosphorylation Regulates Enhancer Activation"

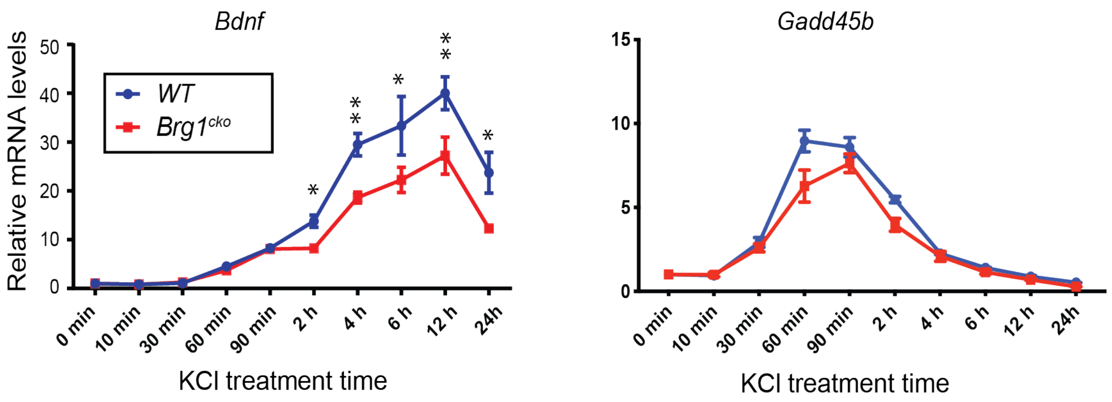

**Figure S1. BRG1 regulates neuronal ARG activation.** Shown are relative mRNA levels of *Bdnf* and *Gadd45b* in cultured *Brg1*$^{cko}$ (red square) and control (blue dot) cortical neurons at different time points after KCl treatment as measured by RT-qPCR. Significance was determined by Student's t-test. *: p<0.05. **: p<0.01.

**A**

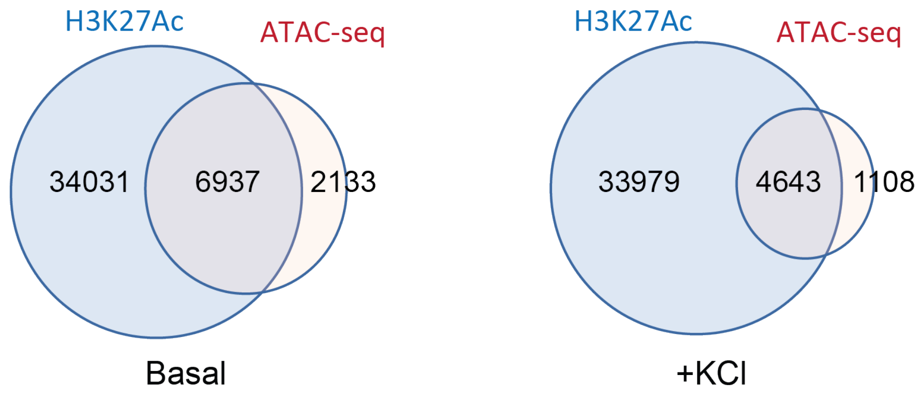

**B**

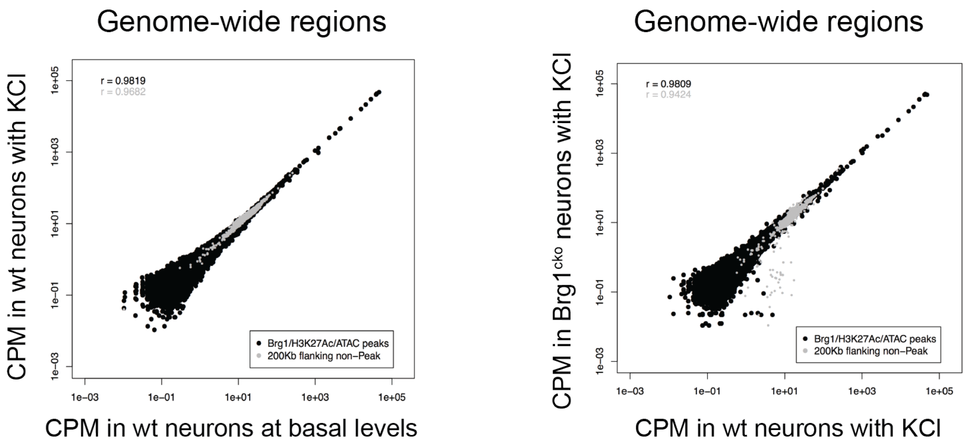

**Figure S2. BRG1 deletion does not change overall chromatin accessibility in neuron cultures.**

A. Overlaps between ATAC-seq peaks and H3K27Ac peaks in cultured wild-type cortical neurons in basal and depolarized conditions.
B. Genome-wide comparison of ATAC-seq peak intensities in wild-type neurons in basal and depolarized conditions (left) and in wild-type and *Brg1*$^{cko}$ neurons in depolarized conditions (right).

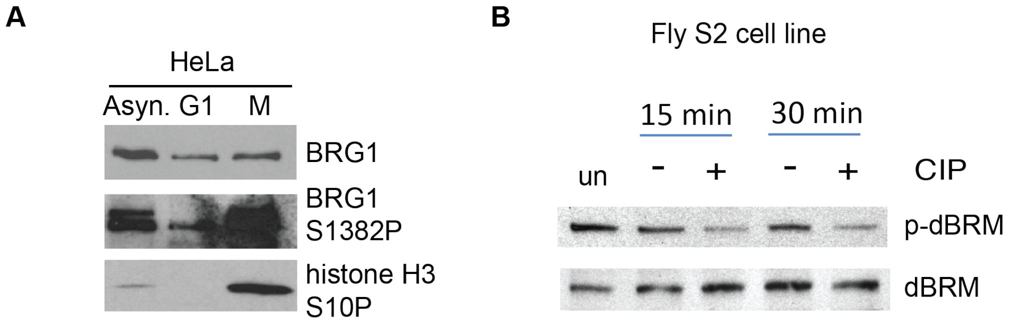

**Figure S3. BRG1 phosphorylation depends on cell-cycle stage, and antibody recognizes *Drosophila* BRM in S2 cells.**

A. HeLa cells were synchronized and arrested at G1 or M phase. Lysates were blotted with indicated antibodies.

B. *Drosophila* S2 cell lysates treated with or without CIP were blotted with antibodies against BRG1 and phosphorylated BRG1, both of which cross-react to *Drosophila* BRM.

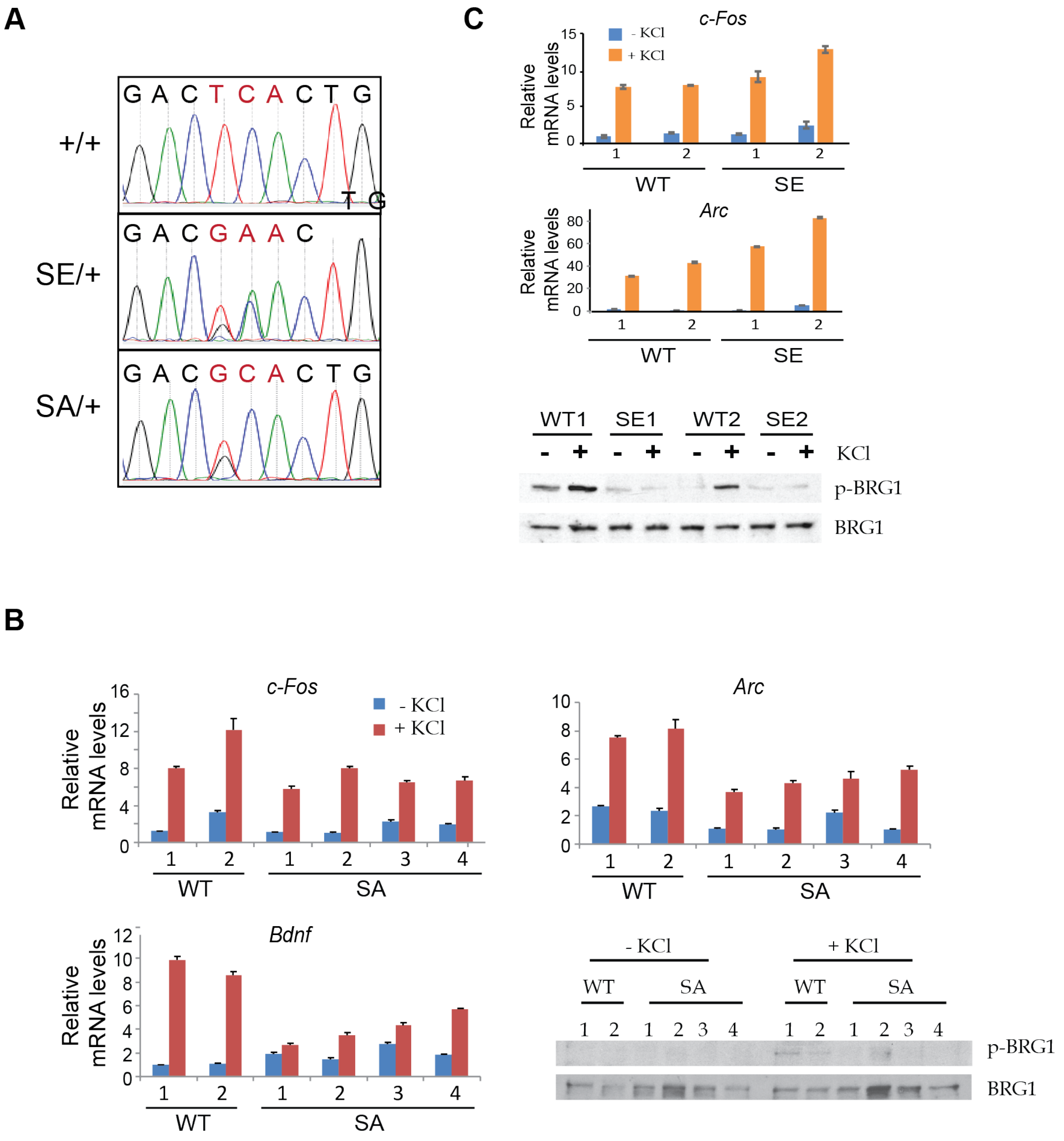

**Figure S4. Generation and characterization of Brg1-SA and Brg1-SE neurons.**

A. Sequencing of relevant genomic regions of *SA* and *SE* mice demonstrate that desired mutations were introduced in the region of BRG1 S1382. The changed codons are in red.
B. mRNA levels of *c-Fos* and *Arc* in wild-type and *SA* neurons with or without 5-h KCl stimulation. Numbered are neurons from different embryos. Western blots show BRG1 levels and the specificity of phospho-specific BRG1 antibody.
C. mRNA levels of ARGs in wild-type and *SE* neurons with or without 5-h KCl stimulation. Numbered are neurons from different embryos. Western blots show BRG1 levels and the specificity of phospho-specific BRG1 antibody.

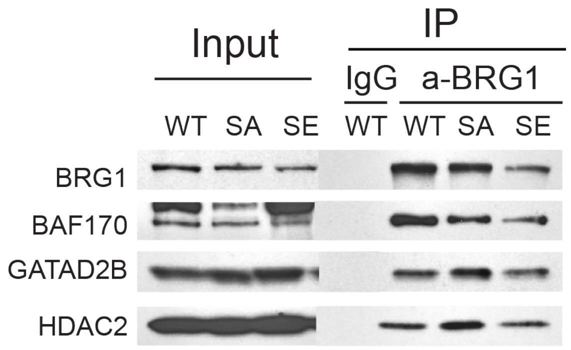

**Figure S5. Differential interactions of NuRD subunits with BRG1 phospho-mutants.** Western blot analyses of samples immunoprecipitated from nuclear extracts from wild-type and *SA* or *SE* P5 cortices using antibodies against BRG1. BAF170 is a control for BAF subunits.

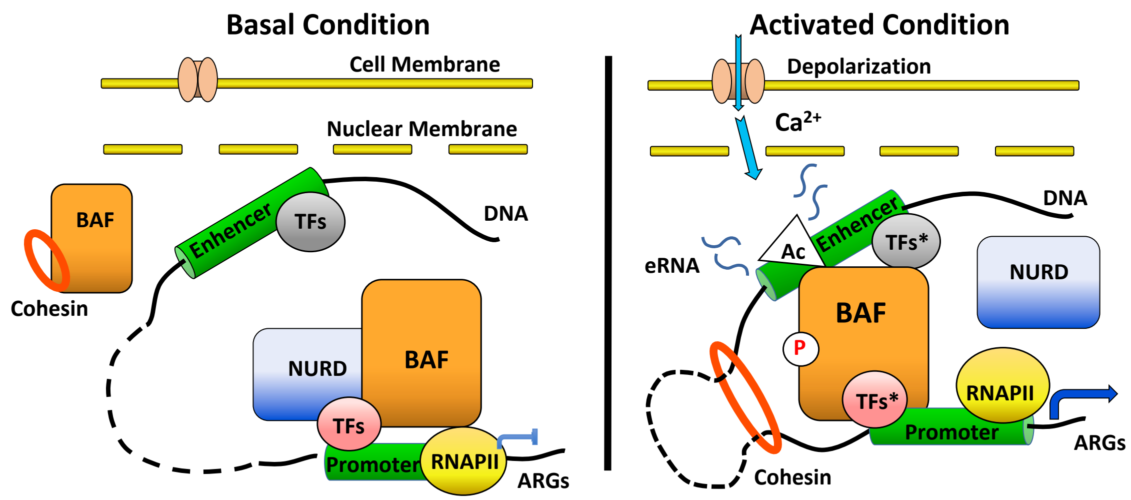

**Figure S6. Model of BRG1 phosphorylation in enhancer activation.** At basal condition, the basal binding of unphosphorylated BRG1, together with the repressive NuRD complex, prevent the activation of enhancers and the expression of ARGs. There is also low level of cohesin at inactive enhancers and no enhancer-promoter looping. Upon neuronal stimulation and activation of $Ca^{2+}$ signaling, initially increased H3K27Ac and activated transcription factors (TFs*) at enhancers recruit more BRG1/BAF to enhancers. $Ca^{2+}$ signal induced BRG1 phosphorylation leads to the local dissociation of repressors such as the NuRD complex, as well as an increase of cohesin mediated enhancer-promoter lopping, possibly through enhanced cohesin dynamics. Together with activated transcription factors and co-factors, these events lead to further increased H3K27Ac and RNA pol II recruitment, expression of eRNAs and ultimately ARG mRNA expression.

**Table S1. Comparison of mass spec peptide numbers and coverages of BAF subunits and key interacting proteins.**

| | Protein | Bassal | | Depolarized | | Depol./Basal (#) | Depol./Basal (%) |
|---|---|---|---|---|---|---|---|
| | | peptide # | coverage % | peptide # | coverage % | | |
| Main BAF subunits | BRG1 | 277 | 32.9 | 270 | 30.5 | 0.97 | 0.93 |
| | BRM | 250 | 34 | 225 | 30.3 | 0.90 | 0.89 |
| | ARID1A | 77 | 20.1 | 60 | 16.1 | 0.78 | 0.80 |
| | BAF170 | 176 | 29 | 206 | 30.9 | 1.17 | 1.07 |
| | BAF155 | 108 | 26.5 | 97 | 23 | 0.90 | 0.87 |
| | BAF47 | 30 | 21 | 37 | 23.1 | 1.23 | 1.10 |
| | BAF57 | 59 | 41.8 | 68 | 36.3 | 1.15 | 0.87 |
| | BAF53a | 26 | 26.1 | 18 | 22.6 | 0.69 | 0.87 |
| | BAF53b | 45 | 31.9 | 36 | 32.6 | 0.80 | 1.02 |
| | BAF60a | 67 | 35.5 | 62 | 35.9 | 0.93 | 1.01 |
| | BAF60c | 66 | 38.5 | 71 | 40.2 | 1.08 | 1.04 |
| | CREST | 14 | 12.7 | 19 | 15.2 | 1.36 | 1.20 |
| | PBRM1 | 56 | 18.8 | 37 | 16.6 | 0.66 | 0.88 |
| | BRD7 | 31 | 32 | 26 | 25 | 0.84 | 0.78 |
| | BRD9 | 5 | 9.2 | 7 | 6.4 | 1.40 | 0.70 |
| | 45a/PHF10 | 6 | 6.2 | 6 | 10.1 | 1.00 | 1.63 |
| | 45b/DPF1 | 41 | 35.7 | 35 | 27.4 | 0.85 | 0.77 |
| | 45c/DPF3 | 15 | 16.9 | 12 | 11.9 | 0.80 | 0.70 |
| | 45d/DPF2 | 23 | 30.2 | 19 | 29.4 | 0.83 | 0.97 |
| | ACTB | 86 | 44 | 86 | 45.6 | 1.00 | 1.04 |
| Kinase | CaMK2B | 12 | 12.2 | 9 | 12.5 | 0.75 | 1.02 |
| Repressor (NuRD subunits) | GATAD2B | 25 | 32.8 | 15 | 16.7 | 0.60 | 0.51 |
| | MTA3 | 11 | 10.7 | 7 | 5.6 | 0.64 | 0.52 |
| | MTA1 | 10 | 11.3 | 7 | 7.7 | 0.70 | 0.68 |
| | HDAC2 | 17 | 15.4 | 11 | 18.9 | 0.65 | 1.23 |
| | CHD4 | 8 | 3.9 | 0 | 0 | 0.00 | 0.00 |
| Cohesin | SMC3 | 4 | 3.5 | 1 | 1 | 0.25 | 0.29 |

**Table S2. Primers used in the study.**

**RT-qPCR**

| | |
|---|---|
| mBdnf F | CAGTACAGTGGTTCTACAATC |
| mBdnf R | CAATTCACAATTAAAGCAGC |

| | |
|---|---|
| Gadd45b coding F | GTTCTGCTGCGACAATGACA |
| Gadd45b coding R | TTGGCTTTTCCAGGAATCTG |

| | |
|---|---|
| *c-fos* coding(F) | ATCCTTGGAGCCAGTCAAGA |
| *c-fos* coding(R) | ATGATGCCGGAAACAAGAAG |

| | |
|---|---|
| *c-fos* e1(F) | TCCGGTAAGGGCATTGTAAG |
| *c-fos* e1(R) | CAAAGCCAGACCCTCATGTT |
| *c-fos* e2(F) | TGCAGCTCTGCTCCTACTGA |
| *c-fos* e2(R) | GAGGAGCAAGACTCCCACAG |
| *c-fos* e3(F) | GGGGTGGTGCTAGTGGTAAA |
| *c-fos* e3(R) | TCATCGTATGCCTCTGCTTG |
| *c-fos* e4(F) | AGGGGGACACATGAGTTCTG |
| *c-fos* e4(R) | GACAAGCCGGGTTAGACTGA |
| *c-fos* e5(F) | CAACCCTGTCATCTATTTAGC |
| *c-fos* e5(R) | TAAGAACTGCGGGGGTCTTC |

| | |
|---|---|
| Arc coding forward | GTGAAGACAAGCCAGCATGA |
| Arc coding reverse | CCAAGAGGACCAAGGGTACA |

| | |
|---|---|
| Arc enhancer forward | GGCTGGAGACTGGTGACATT |
| Arc enhancer reverse | CCATCTGCTTTCTCCTGGAA |

| | |
|---|---|
| mNr4a1 F | GTTATCCGAAAGTGGGCAGA |
| mNr4a1 R | AGTACCAGGCCTGAGCAGAA |

| | |
|---|---|
| mKcna1 F | AGGCTCAGTTGCTCCATGTT |
| mKcna1 R | TCAGCTGTGGTGCAGTTACC |

| | |
|---|---|
| Egr1 F | AGACGAGTTATCCCAGCCAAA |
| Egr1 R | GGTCGGAGGATTGGTCATGC |

| | |
|---|---|
| mJunb F | TTTTGTCAAAGCCCTGGACG |
| mJunb R | GGGGAGTAACTGCTGAGGTT |

**ChIP_qPCR**

FosP_ChIP1
F: GTGACGTAGGAAGTCCATCCA
R: CCCGAGAACATCATGGTCGA

Fos e2_ChIP
F: TGCAGCTCTGCTCCTACTGA
R: GAGGAGCAAGACTCCCACAG

ArcP_ChIP
F: GAGGAGCTTAGCGAGTGTGG
R: AGCATAAATAGCCGCTGGTG

Arc e_ChIP
F: GGCTGGAGACTGGTGACATT
R: CCATCTGCTTTCTCCTGGAA

Bdnf ChIP
F: AGCCCTCCAGAACCTAGTCA
R: CAGAAGCCAAGCCTGTCATTA

Cd4 ChIP
F: GCAAGATAGCTAAGCCAAACACATT
R: CACCCTACGCTGACATAGTGGTTC

**3C**

c-Fos pro-rev anchor primer CTGACGGATAAACATTGTGC

c-Fos-A primer CATTCTTGGTTTCCACAGGC

c-Fos-B primer GATATTAGAGTTAAGGTGTGC

c-Fos-C primer CATCTTTGACCTATCAGCAC

c-Fos-D primer CAAGCTTGAACAGTTAGGTAG

Arc pro-rev anchor primer AGTCAGTTGAGGCTCAGCAA

Arc-A primer GAGGTCTTGTATCCTGGCTG

Arc-B primer CACACCAAATCTGCAGAGATT

Arc-C primer GGCTGGAGACTGGTGACATT

Arc-D primer CAGGTCCTGTCTATGCTCTT
